# Splanchnic and pelvic spinal afferent pathways relay sensory information from the mouse colorectum into distinct brainstem circuits

**DOI:** 10.1101/2025.02.24.639793

**Authors:** QingQing Wang, Alice E. McGovern, Melinda Kyloh, Grigori Rychkov, Nick J. Spencer, Stuart B. Mazzone, Stuart M. Brierley, Andrea M. Harrington

**Author notes:** **Correspondence:** Dr. Andrea M. Harrington.

## Abstract

This study aimed to identify where the sensory information relayed by the two spinal afferent pathways innervating the distal colon and rectum (colorectum), the splanchnic and pelvic spinal afferent pathways, integrates within the brainstem. Localised injections of transneuronal viral tracer (herpes simplex virus H129 strain expressing EGFP (H129-EGFP)) into the distal colon was used to assess the brainstem structures receiving ascending input from the colorectum. H129-EGFP+ cells were distributed in structures involved in ascending sensory relay, descending pain modulation and autonomic regulation in the medulla from 96 hours and in pontine and caudal midbrain 120 hours after inoculation. In a separate cohort of mice, *in vivo* noxious colorectal distension (CRD) followed by brainstem immunolabelling for phosphorylated MAP kinase ERK 1/2 (pERK) showed that many of the structures in which H129-EGFP+ labelling was observed were relevant to colorectal sensory processing. Surgical removal of dorsal root ganglia (DRG) containing cell bodies of splanchnic colorectal afferent neurons, significantly reduced CRD evoked neuronal activation within the caudal ventrolateral medulla, rostral ventromedial medulla and the lateral parabrachial nuclei. Whilst, removal of DRG containing cell bodies of pelvic colorectal afferent neurons significantly reduced CRD evoked neuronal activation within the rostral ventromedial medulla, lateral parabrachial nuclei, the locus coeruleus, Barrington’s nucleus and periaqueductal gray. Collectively, this study showed that the two spinal afferent pathways innervating the colorectum differentially shape colorectal processing within the brainstem and provides new insight into their unique roles to mediating visceromotor responses and defecation associated with colorectal nociception.

## 2 Introduction

Sensory information from the distal colon and the rectum (colorectum) is conveyed into the brain primarily via spinal ascending pathways (Westlund 2000; Sikandar *et al*. 2012; Kyloh *et al*. 2022; Harrington 2023). The colorectum is innervated by two spinal afferent pathways, the lumbar splanchnic and the sacral pelvic (Osman *et al*. 2023; Wang *et al*. 2024). The unique roles of these two spinal afferent nerves in shaping where colorectal sensory information is relayed to within the brain are not fully known. Splanchnic colorectal afferent fibres relay information into the thoracolumbar (TL, T10-L1) spinal cord, while colorectal pelvic afferent fibres project into the lumbosacral (LS, L5-S1) spinal cord (Harrington *et al*. 2019; Wang *et al*. 2024). Splanchnic and pelvic colorectal afferent fibres possess different mechanosensitivity profiles, with a large proportion of colorectal splanchnic afferent fibres being high-threshold nociceptors, whilst pelvic afferent fibres are more functionally diverse (Brierley *et al*. 2004). Correspondingly, our recent studies in mice show that the dorsal horn circuits in the TL spinal cord processing colorectal afferent input are relevant to the relay of nociceptive information into the brain, whilst those in the LS spinal cord demonstrate elements of supraspinal and intraspinal transmission of nociceptive and wide dynamic information, in addition to supporting parasympathetic motor reflexes (Harrington *et al*. 2019; Wang *et al*. 2024). This study aimed to determine if these differences extend further along the ascending spinal pathway and assessed how the ascending output from the TL and LS spinal cord shapes colorectal sensory processing within the brainstem.

The brainstem is the first site at which viscerosensory information is integrated within the brain to support behavioural and affective motor responses associated with colorectal nociception (Westlund 2000). This involves brainstem structures caudal to the midbrain, with decerebration at the level of the brainstem pontine, but not rostral to the brainstem midbrain, attenuating the visceromotor responses evoked by noxious colorectal distension (CRD) (Ness *et al*. 1988). Spinal cord transection (at T8) significantly attenuates transneuronal labelling of cells from the distal colon within pontine and midbrain structures and partially within the caudal medulla reticulum (Vizzard *et al*. 2000). Consistent with this, several of the brainstem structures identified by distribution mapping of neuronal activation evoked by colorectal distension (CRD) (Monnikes *et al*. 1994; Monnikes *et al*. 2003; Rouzade-Dominguez *et al*. 2003) and transneuronal viral tracing from the colorectum (Pavcovich *et al*. 1998; Valentino *et al*. 2000; Vizzard *et al*. 2000; Rouzade-Dominguez *et al*. 2003; He *et al*. 2018) are known targets of ascending output from the spinal cord (Wang *et al*. 1999; Polgar *et al*. 2010; Todd 2010; Martins *et al*. 2017; Wercberger *et al*. 2019). Many of these structures influence pain transmission and autonomic outflow from the spinal cord via their extensive output into brainstem-wide networks or descending projections into the spinal cord (Ness *et al*. 1988; Monnikes *et al*. 1994; Ness *et al*. 1998; Pavcovich *et al*. 1998; Naitou *et al*. 2018; Nakamori *et al*. 2018; Lyubashina *et al*. 2019; Nakamori *et al*. 2019; Lyubashina *et al*. 2022).

Our first aim was to identify the brainstem structures connected to the spinal afferent pathways ascending from the colorectum, given studies mapping these circuits in mice are lacking. To do this, we injected a Herpes simplex virus type 1 strain (H129) expressing EGFP (HSV-H129-EGFP) that has a predominant anterograde mode of transmission into the colorectal wall (Rinaman *et al*. 2004). Injections were localised to the region of the colorectum that we have shown previously to be densely innervated by splanchnic and pelvic spinal afferent fibres, but sparsely by vagal afferent fibres (Wang *et al*. 2024). The distribution pattern of H129-EGFP labelling was then tracked through the spinal cord and within the brainstem in a time-dependent manner. *In vivo* noxious colorectal distension (CRD) followed by immunolabelling for neuronal activation marker phosphorylated MAP kinase ERK 1/2 (pERK) was used to establish the brainstem nuclei functionally relevant to colorectal sensory processing. We then aimed to determine the relative contribution of signalling via the TL and LS spinal cord on colorectal processing within brainstem structures relevant to i) ascending relay, ii) descending pain modulation and iii) autonomic outflow. To do this, we used a dorsal root ganglion (DRG) removal model to assess the effect of attenuating colorectal signaling via either the TL or LS spinal cord on CRD evoked neuronal activation within the brainstem. We have previously shown in this DRG removal model (Kyloh *et al*. 2022), that visceromotor responses evoked from rectal distension, at pressures in the physiological and noxious ranges, are attenuated by removing LS DRG (L5-S1) containing the cell bodies of colorectal pelvic afferents. The visceromotor responses evoked by noxious distension of the distal colon were attenuated by removing the TL DRG (T13-L1), which contain the cell bodies of colorectal splanchnic afferents. The outcomes of these assessments provided direct evidence of the unique roles splanchnic and pelvic spinal afferent pathways play in shaping colorectal sensory processing in the brainstem.

## 3 Materials and methods

### 3.1 Animals

Female and male C57BL/6J (8-16 weeks, weight range 18-30 grams) mice were used for all methods. All animal procedures were performed in accordance with the approval of Animal Ethics Committees of the South Australian Health and Medical Research Institute (SAHMRI; Application SAM190), The University of Melbourne (Application 1714311.1) and Flinders University (Approval no.861-13). The HSV1 H129-EGFP transneuronal tracing and pERK neuronal activation studies were performed in separate cohorts of mice.

### 3.2 HSV1 H129-EGFP transneuronal tracing from the colorectal wall

Anterograde transneuronal tracing was performed in female and male mice from the colorectum using a herpes simplex virus 1, H129 strain, expressing EGFP (HSV1 H129-EGFP). The wildtype HSV-1 H129 origin, construction of the H129-EGFP recombinant virus and its validation for preferential trans-synaptic movement in the anterograde direction to label the central circuitry arising from peripheral structures has previously been described (Barnett *et al*. 1995; Rinaman *et al*. 2004; McGovern *et al*. 2012; McGovern *et al*. 2012; McGovern *et al*. 2015; Wojaczynski *et al*. 2015). Aliquots of viral stocks were thawed from −80◦C and 10 μl of 7 ×10^7^ pfu/ml of H129-EGFP virus was injected into the subserosa-musculature wall of the colorectum (covering 1 cm proximal to the pelvic bone through to immediately below the pelvic bone. Within this region, 3-6 injections (1-2 μl/injection) were made across bilateral and midline locations **(Fig. 1Ai)** using a 30-gauge needle (HAMC7803-07, point style: 4 (10-12°); Hamilton Company, Bio-Strategy, Campbellfield, Vic, Australia) attached to Hamilton 5 μl syringe (HAMC7634-01 5 µl 700 series RN syringe; Hamilton Company, Bio-Strategy). The needle tip was inserted into the sub-serosal wall space and tunnelled a short distance caudally, ensuring the needle tip was always visible within the wall. The viral solution was expelled as the needle was gradually pulled out of the needle track. Care was taken to minimize leakage of the virus during injections into the colon wall, by using cotton tip applicators at injection sites. The abdominal incision was then sutured closed and following recovery, mice were group-housed and monitored twice daily for signs of weight loss, altered colonic motility (changes to fecal output and form) and stereotypical behaviour (altered gait or locomotion, reduced or excessive grooming, lack of nesting, repetitive movements). Mice showed no clinical signs 24-48 hours post-viral tracing and in varying severity from 72 hours through to 120 hours post-inoculation. Mice underwent transcardial perfuse fixation at designated survival points or upon accumulative or severe clinical signs outlined above. Tissue (DRG, spinal cord and brain) was collected 24 hours post-inoculation (N=3 mice inoculated, 2F:1M), 48 hours post-inoculation (N=4 mice inoculated, 2F:2M), 72 hours post-inoculation (N=4 mice inoculated, 2F:2M), 96 hours post-inoculation (N=5 mice inoculated, 3F:2M) and 120 hours post-inoculation (N=10 mice inoculated, 7F:3M).

**Figure 1:**
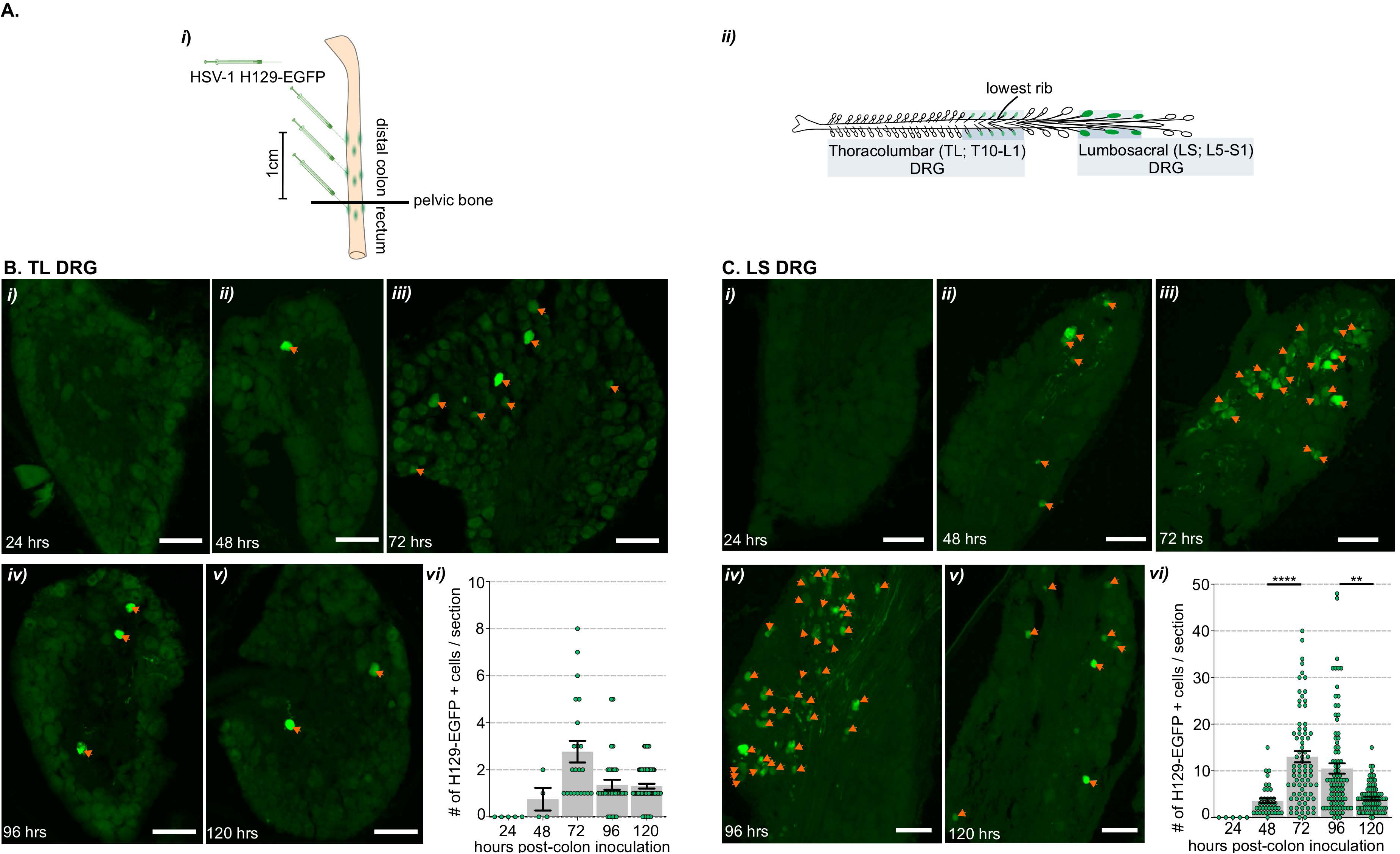
Distribution of H129-EGFP+ cells within the thoracolumbar (TL) and lumbosacral (LS) dorsal root ganglia following colorectal HSV-1 H129-EGFP inoculation. **A)** Schematic showing **(i)** the injection sites of HSV-1 H129-EGFP (green) at multiple sites into the subserosa-musculature wall of the mouse distal colon and rectum (colorectum), covering the region 0.2 cm below the pelvic bone through to 1 cm proximal to the pelvic bone and **(ii)** the dorsal root ganglia (DRG) collected at spinal levels thoracic 10 through to lumbar 1 (Thoracolumbar, TL. T10-L1) and lumbar 5 through to sacral 1 (Lumbosacral, LS. L5-S1) in which HSV-1 H129-EGFP labelling was assessed in 24-hour increments after colorectal inoculation. **(B-C)** Representative photomicrographs of H129-EGFP+ (green, indicated by orange arrows) cells within sections of **B)** TL DRG (T10-L1) and **C)** LS DRG (L5-S1) at **(i)** 24-, **(ii)** 48-, **(iii)** 72-, **(iv)** 96- and **(v)** 120-hours post-inoculation of H129-EGFP+ within the colorectal wall. Scale bars: 100 μm. **(vi)** Quantification of the total number (#) of H-129 EGFP+ cells within sections of DRG. \*\**P*<0.01 and \*\*\*\**P*<0.0001 determined by a One-way Kruskal-Wallis test (non-parametric data), comparisons were made at every time point. Data points represent the individual DRG sections, gained from 2-10 sections/mouse, collected 24 hours (N=3 mice 2F:1M), 48 hours (N=4 mice 2F:2M), 72 hours (N=4 mice, 2F:2M), 96 hours (N=3 mice 1F:2M) and 120 hours (N=3 2F:1M) post-HSV-1 H129-EGFP inoculation. Scale bars: 100μm.

### 3.3 *In vivo* Colorectal Distension (CRD)

Mice (N=10 mice, 5F:5M) separate to those that underwent HSV1 H129-EGFP transneuronal tracing underwent *in vivo* CRD in order to identify neurons activated by colorectal input. Following isoflurane anaesthesia, mice were given an enema of sterile saline followed by insertion of a 2 cm balloon catheter into the perianal canal. When the base of the balloon was localised approximately 0.2 cm from the anal verge, the catheter tube was secured in place by taping it to the base of the tail (Christianson *et al*. 2007). Mice were then placed in a Perspex box, the balloon catheter was distended to a pressure of 80 mmHg applied via a syringe attached to three-way tap and monitored via a sphygmomanometer. Distension was held for 10 seconds, then released for 5 seconds. This sequence was repeated 5 times (Harrington *et al*. 2019; Wang *et al*. 2024). Immediately after the fifth distension, mice were placed into an isoflurane induction chamber and, within 30-40 seconds, given an overdose of euthanasia agent (Lethabarb via intraperitoneal injection) and underwent transcardial perfusion fixation. Fixation was completed within 5-6 minutes after the final distension. A separate group of mice (N=10 mice, 5F:5M) underwent balloon insertion without distension (no CRD).

### 3.4 Surgical removal of TL and LS DRG

In order to compare how signaling via the splanchnic or pelvic afferent pathways shape colorectal evoked neuronal activation in the brainstem, a separate cohort of female mice underwent surgical removel of DRG at either vertebrae levels T13-L1 (thoracolumbar DRG TL removed) or L5-S1 (lumbosacral DRG LS removed) (Kyloh *et al*. 2022) or sham surgery 8-9 days prior to *in vivo* CRD procedure described above (N=5 mice/group). Mice underwent aneasthesia induction using inhalation isoflurane at 4% in 1L/min oxygen which was then maintained at 1.5-2% in 1L/min oxygen. Animals were positioned on a thermostat-controlled heat mat to maintain body temp throughout the procedure (Adloheat, Pakenham, Vic, Australia). Before incision, animals were administered via subcutaneous (s.c) injection, 0.5mg/kg buprenorphine (Temvet). The dorsal surface was shaved and cleaned with 0.5% chlorhexidine and 70% alcohol swab (Briemar). An incision (∼20mm in length) was made along the dorsal midline and skeletal muscles retracted to expose the vertebral column. A laminotomy was performed to remove small fragments of vertebral bone from the dorsal (uppermost) surface of each DRG to expose the ganglion but not the dura or spinal cord. The dorsal nerve root between DRG located at both sides of the vertebrae at levels T13-L1 (thoracolumbar DRG TL removed) and L5, L6 and S1 (lumbosacral DRG LS removed) was severed on either side of the ganglion and the ganglion was removed entirely (Kyloh *et al*. 2022). Sham experimental groups underwent laminotomy and DRG exposure but no DRG removal surgery. Following removal of DRG or sham surgery, the wound was irrigated with 0.5% Bupivicaine (Marcain, AstraZeneca) and the muscle closed with individual 5.0 polyglycholic acid absorbable suture (Silverglide). Skin was closed with 6.0 Nylon non-absorbable suture (Silverglide) and site cleaned with 0.5% chlorhexidine and 70% alcohol swab. Prior to withdrawal of anaesthesia, animals were administered a second s.c dose of 0.5mg/kg buprenorphine (Temvet) and s.c antibiotics, 100mg/kg Ampicillin (Alphapharm) and 10mg/kg Baytril (Bayer). Following withdrawal of anaesthesia animals recovered on a heat mat until fully mobile and then returned to their home cage. Postoperatively, animals received 0.1mg/kg oral buprenorphine (Schering Plough) in Nutella (Ferrero) paste at 24-hour intervals for 72 hours.

### 3.5 Transcardial perfuse fixation and tissue processing

Mice were euthanised with an overdose of Lethabarb (60 mg/kg pentobarbitone sodium solution, Virbac Australia, Milperra, NSW, Australia), via intraperitoneal injection, prior to opening the chest cavity and 0.5ml heparin-saline was then injected into the left ventricle followed by insertion of a 22-gauge needle, attached to a peristaltic perfusion pump. The right atrium was then snipped allowing for perfusate drainage. Warm saline (0.85 % physiological sterile saline) was perfused prior to ice-cold 4% paraformaldehyde in 0.1 M phosphate buffer (Sigma-Aldrich, MO, USA). Following complete perfusion, spinal cord and brain, and in the case of HSV1-H129-EGFP inoculated mice, thoracolumbar T10-L1 and lumbosacral L5-S1 DRG, were collected (Robinson *et al*. 2004; Christianson *et al*. 2006). The lowest rib was used as an anatomical marker of T13 DRG. The spinal cord levels T10-L1 were identified by DRG root insertion points or vertebra level and the spinal cord from below vertebra level L2 was removed, which contains spinal cord levels L5-S1. All tissue was post-fixed in 4% paraformaldehyde in 0.1 M phosphate buffer at 4^°^C for 18-20 hours and then cryoprotected in 30% sucrose/phosphate buffer (Sigma-Aldrich) overnight at 4^°^C. Ganglia were then placed in 100% OCT (Tissue-Tek^®^ O.C.T. Compound, Sakura^®^ Finetek, Netherlands) and snap-frozen in liquid nitrogen. The spinal cord and brain underwent an additional 24 hour incubation at 4^°^C in 50% OCT/30% sucrose/phosphate buffer solution before freezing in 100% OCT. Tissue was cryosectioned and sections were placed onto gelatin coated slides for visualization of H129-EGFP-labelling (DRG, spinal cord and brain 20-50 μm sections) or for immunolabelling (spinal cord and brain 10 μm thick sections). Sections for H129-EGFP-labelling visualization were air dried for an hour before being washed in 0.2% Triton-X 100 (Sigma-Aldrich) in 0.1M phosphate buffered saline before cover slipping with ProLong Diamond Antifade Mountant with DAPI (P36966; Invitrogen, ThermoFisher Scientific). For immunolabelling, two slides (100 μm apart) covering medulla, pons, midbrain levels were randomly selected per mouse. The spinal cord and brain levels were identified *ex vivo* using the Allen Institute Mouse Spinal Cord Atlas (https://mousespinal.brain-map.org/imageseries/showref.html) (Lein *et al*. 2007) and the Paxinos and Franklin Mouse Brain Atlas (Paxinos *et al*. 2001).

### 3.6 pERK immunolabelling

Immunolabelling for pERK using HRP-DAB detection (EnVision FLEX Mini Kit, Agilent) and the DAKO Omnis auto-stainer (Agilent Technologies Australia, Mulgrave, Australia), followed by hematoxylin staining, was performed as previously described (Wang *et al*. 2024)using the primary antibody Rabbit anti-pERK ½ (MAB4370, Cell Signalling Technology, Genesearch, Qld). Rabbit monoclonal antibody detects endogenous levels of p44 and p42 MAP Kinase (Erk1 and Erk2) when dually phosphorylated at Thr202 and Tyr204 of Erk1 (Thr185 and Tyr187 of Erk2), and singly phosphorylated at Thr202. The antibody does not cross-react with the corresponding phosphorylated residues of either JNK/SAPK or p38 MAP kinases (manufacturer’s specifications) and binds specifically to a 44kDa band in stimulated mouse tissue (Miyaji *et al*. 2009).

### 3.7 pERK and neurochemical marker immunolabelling

Immunofluorescence for pERK and neurochemical markers was performed as previously described (Harrington *et al*. 2019; Wang *et al*. 2024). Briefly, after air drying for 1 hour, sections were washed with 0.5% Triton-TX 100 (Sigma-Aldrich, MO, USA) in 0.1M phosphate-buffered saline (T-PBS) to remove excess OCT. Non-specific binding of secondary antibodies where blocked using either 5% normal chicken serum (CHBX0010; Applied Biological Products, SA, Australia) or 5% normal goat serum (GTBX0050; Applied Biological Products) diluted in T-PBS for 30 minutes at room temperature. Tissue sections were then incubated with primary antisera Mouse anti Tyrosine Hydroxylase (TH; Isotype IgG1; 1:1000, Catalogue #: 22941, RRID: AB 572268, ImmunoStar. Inc, Hudson, WI, USA) or Goat anti serotonin (5HT; 1:1000, Catalogue #: 20079, RRID: AB_572262, ImmunoStar. Inc) for 24 hours at room temperature, diluted in T-PBS. Sections were then washed in T-PBS and incubated for 1 hour at room temperature with appropriate secondary antibody conjugated, either Chicken anti-rabbit IgG1 AF488 (1:200, Catalogue #: A-21441, RRID: AB_ 2535859), Chicken anti-mouse IgG1 AF594 (1:200, Catalogue #: A-21125, RRID: AB_141593, ThermoFisher Scientific) or Chicken anti-goat AF594 (1:200, Catalogue #: A-21468, RRID: AB_141859, ThermoFisher Scientific). Sections were then washed in T-PBS before mounting in ProLong Glass Antifade Mountant with NucBlue and coverslipped. Slides were allowed to dry for 24 hours prior to visualization. The anti-TH and anti-5HT antisera have been used extensively to label regions of the mouse brain that are known to contain TH (Asmus *et al*. 2008; Churchill *et al*. 2019) and 5HT (Iwasaki *et al*. 2018; Hingorani *et al*. 2022) neuronal cell bodies.

### 3.8 Microscopy

Fluorescence was manually visualized using a confocal laser scanning microscope (Leica TCS SP8X, Germany) or autoscanned using an epifluorescence ZEISS Axioscan 7 Slide Scanner (Carl Zeiss Microscopy, Germany) with a 40X air objective (EGFP in spinal cord and brainstem sections only). pERK-DAB staining was imaged using a NanoZoomer slide scanner (Hamamatsu, Japan) with a 40X objective. Confocal images (1024 × 1024 pixels) were obtained with oil immersion 20X-63X objectives and software zoom of 2-4X magnification. Sequential scanning (5-line average) was performed with the following settings using a tunable white light laser and photomultiplier detectors: 495 nm-excitation and 503 / 538 nm-emission detection for EGFP/AF488 and 561 nm-excitation and 570-625 nm-emission detection for AF594. Spinal cord and brainstem sections were optically sectioned (2 µm thick sections), and z-projected images were reconstructed as z-stacks (10-50 µm).

Images were processed using LAS Lite (Leica), ZEN BLUE (Carl Zeiss Microscopy), FIJI (NIH, MD, USA), and CorelDRAW Graphic Suite 2021 (Corel Corporation, CA, USA) software. Other than making moderate adjustments for contrast and brightness, the images were not manipulated in any way. At the time of pERK-DAB image collection, the images produced were assigned random numbers that de-identified their experimental group for subsequent neuronal quantification. The scanned images were then opened and viewed using NDPview2 (Hamamatsu, Japan) software.

### 3.9 Quantification and statistics

Quantification of H129-EGFP-labelled cells was determined from single plane epifluorescence images obtained from the ZEISS auto-scanner, viewed using ZEN BLUE (Zeiss) software, from 10 sections / spinal cord level/mouse/time point and 2-5 sections/brainstem level/mouse/time point. The density of H129-EGFP + cells over time was expressed as scattered (1-10 H129-EGFP cells/section), dense (11-20 H129-EGFP cells/section) and to substantial infections (21+ H129-EGFP cells/section) (Vizzard *et al*. 2000; Bassi *et al*. 2022). Quantification of pERK-immunoreactive neurons was performed from scanned images of pERK-HRP immunolabelled sections, using QuPath (Queens University, Belfast, Northern Ireland) (Bankhead *et al*. 2017). The number of pERK neurons/section was obtained from 5-10 sections / spinal cord region and 1-3 sections/brain level/mouse. Only cells with a neuronal morphological profile and intact nuclei (identified by NucBlue or hematoxylin counter stain) were included in the counts. The data distribution was determined by the GraphPad Prism 9 normality and lognormality Shapiro-Wilk tests. The specific number of mice in each experimental group (N), the samples/sections per mouse (n) and details of the tests used for statistical comparisons between experimental groups are outlined within the relevant figure legends.

## 4 Results

### 4.1 Distribution of HSV-1 H129-EGFP infected cells (H129-EGFP+) in the spinal cord and brainstem at different time points following colorectal inoculation

Following HSV-1 H129-EGFP injection into the colorectal wall **(Fig. 1Ai)**, H129-EGFP+ cells were observed in DRG at vertebrae levels known to contain colorectal sensory afferent neurons **(Fig. 1Aii)**, TL (T10-L1, **Fig. 1Bi-v)** and LS (L5-S1, **Fig. 1Ci-v)**, in varying density from 48 hours through to 120 hours post-inoculation **(Fig. 1Bvi and 1Cvi)**. Within the spinal cord, H129-EGFP+ cells were observed in the dorsal horn at TL **(Fig. 2)** and LS **(Fig. 3)** spinal cord levels in tissue collected at 72, 96 and 120 hours post-inoculation. H129-EGFP+ cells were observed in the medulla **(Fig. 4 and 5)** in tissue collected 96 and 120 hours post-inoculation and in the pons **(Fig. 6)** and caudal midbrain **(Fig. 7)** in tissue collected 120 hours post-inoculation. In the spinal cord and brainstem, H129-EGFP+ cells had varied morphologies, suggesting labelling of glial cells, neuronal cell bodies and their processes, as previously reported (Garcia-Luna *et al*. 2021).

**Figure 2:**
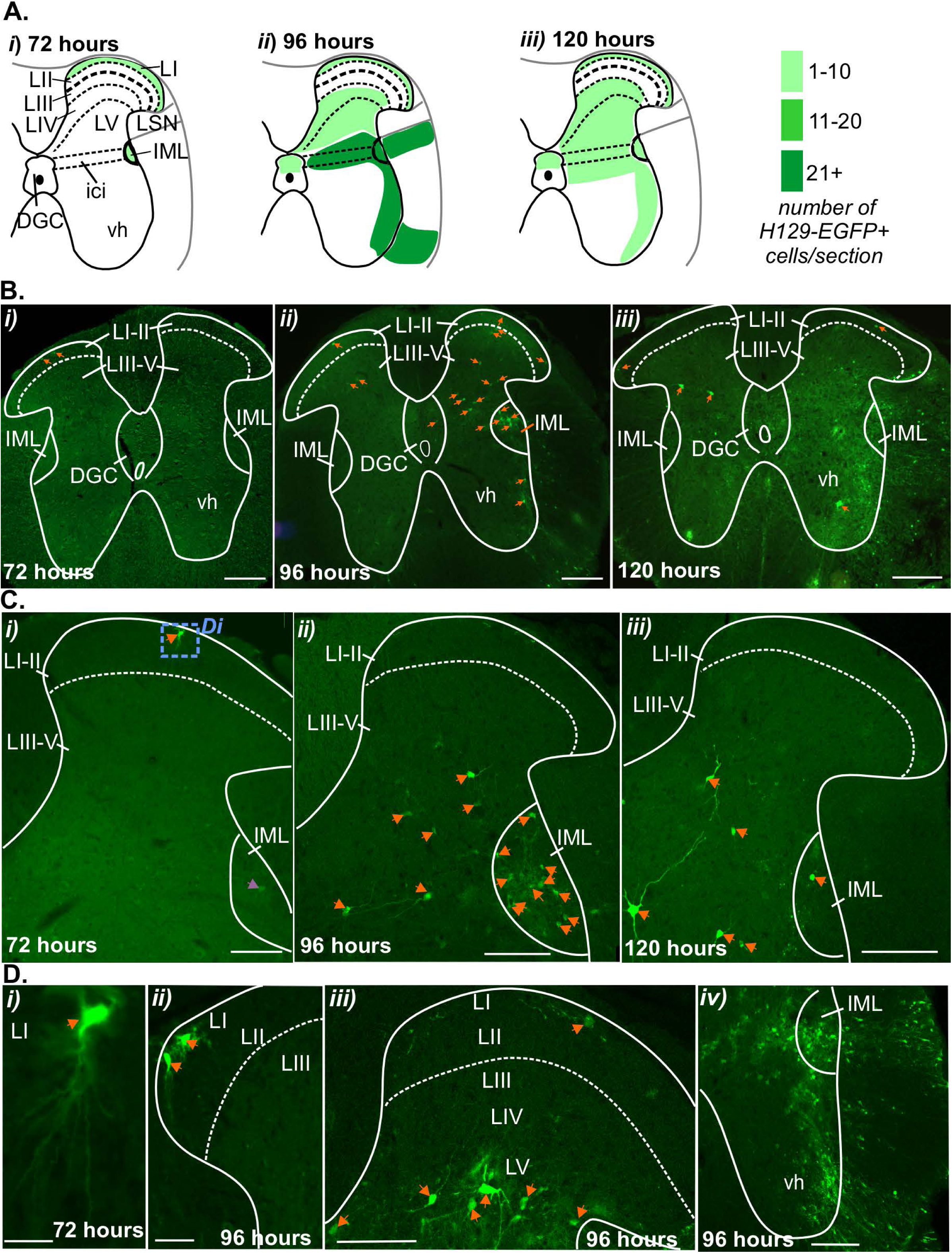
Distribution of H129-EGFP+ cells within the thoracolumbar (TL) spinal cord dorsal horn following colorectal HSV-1 H129-EGFP inoculation. **A**) Schematic (Lein *et al*. 2007) illustration of the TL spinal cord summarizing the distribution (regions indicated by green shading) and semi-quantification of the density (indicated by the intensity of green shading) of H129-EGFP+ cells at 72-, 96- and 120 hours following HSV-1 H129-EGFP inoculation into the colorectum. Data obtained from 5-10 sections/mouse, collected 72 hours (N= 4 mice 2F:2M), 96 hours (N= 5 mice 3F:2M) and 120 hours (N= 8 mice 7F:1M) post-HSV-1 H129-EGFP inoculation. **(B-C)** Representative photomicrographs of the TL spinal cord dorsal horn at **(B)** low magnification and **(C)** high magnification showing the distribution of H129-EGFP+ cells (green, highlighted by orange arrows) in tissue collected at **(i)** 72-, **(ii)** 96- and **(iii)** 120 hours after colorectal inoculation. Scale bars: 100μm. **D)** Photomicrographs of the TL spinal cord at high magnification showing the distribution of EGFP-labelled cells (green, highlighted by orange arrows) within the **(i and ii)** superficial dorsal horn laminae I-II (LI-II), **(iii)** dorsal horn laminae I-V (LI-V) and **(iv)** ventral horn in tissue collected at **(i)** 72- and **(ii-iv)** 96-hours post-inoculation. The image in **Di** (scale bar: 20 μm) corresponds to the region within the blue dotted box in **Ci**. Scale bars: 100 μm. **(B-D)** Dashed lines indicate the border between lamina II (LII) and lamina III (LIII). Abbreviations: LI= lamina I, lamina IV= lamina 4, lamina V= lamina 5, ici= intercalated nuclei and IML= intermediolateral nuclei.

**Figure 3:**
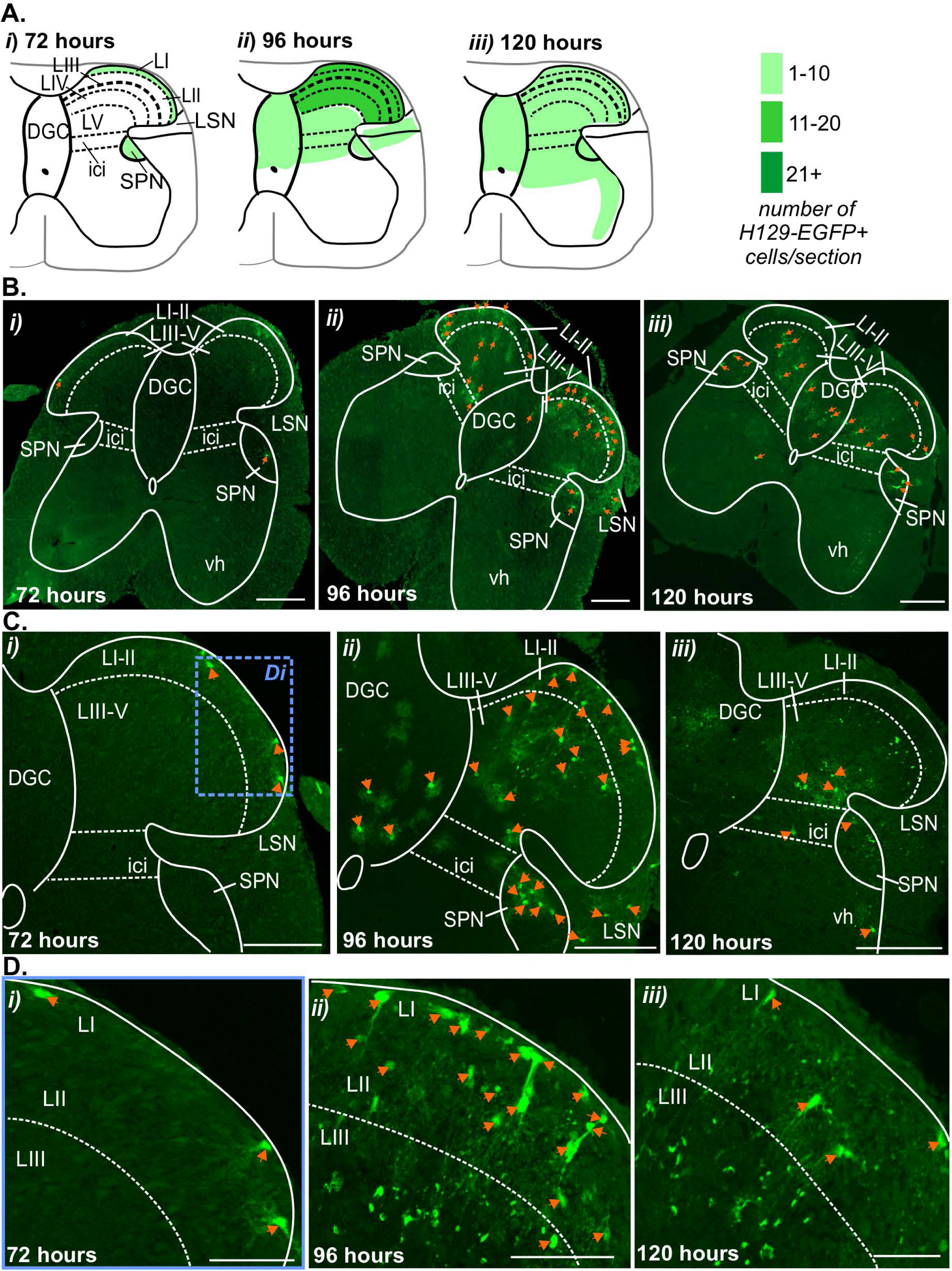
Distribution of H129-EGFP+ cells within the lumbosacral (LS) spinal cord dorsal horn following colorectal HSV-1 H129-EGFP inoculation. **A)** Schematic (Lein *et al*. 2007) illustration of the LS spinal cord summarizing the distribution (regions indicated by green shading) and semi-quantification of the density (indicated by the intensity of green shading) of H129-EGFP+ cells at 72-, 96- and 120 hours following HSV-1 H129-EGFP colorectal inoculation. Data obtained from 5-10 sections/mouse, collected 72 hours (N= 4 mice 2F:2M), 96 hours (N= 5 mice 3F:2M) and 120 hours (N= 8 mice 7F:1M) post-HSV-1 H129-EGFP inoculation. **(B-D)** Representative photomicrographs of the LS spinal cord dorsal horn at **(B)** low magnification and **(C)** high magnification showing the distribution of H129-EGFP+ cells (green, highlighted by orange arrows) in tissue collected at **(i)** 72-, **(ii)** 96- and **(iii)** 120 hours after colorectal inoculation. **D)** Photomicrographs of the LS spinal cord at high magnification showing the distribution of H129-EGFP+ cells (green, highlighted by orange arrow) within the superficial dorsal horn laminae I-III (LI-III), in tissue collected at **(i)** 72-, **(ii)** 96- and **(iii)** 120 hours post-inoculation of HSV-1 H129-EGFP within the colorectal wall. Image **(Di**) corresponds to the region within the blue dotted box in **(Ci)**. Scale bars: **B and C:** 100 μm and **D:** 20 μm. **(B-D)** Dashed lines indicate the border between lamina II (LII) and lamina III (LIII), and the lateral edges of the intercalated nuclei (ici). Abbreviations: DGC= dorsal grey commissure, LI= lamina I, LII= lamina II, LIII= lamina III, lamina IV= lamina 4, lamina V= lamina 5, LSN= lateral spinal nucleus and SPN= sacral parasympathetic nuclei.

**Figure 4:**
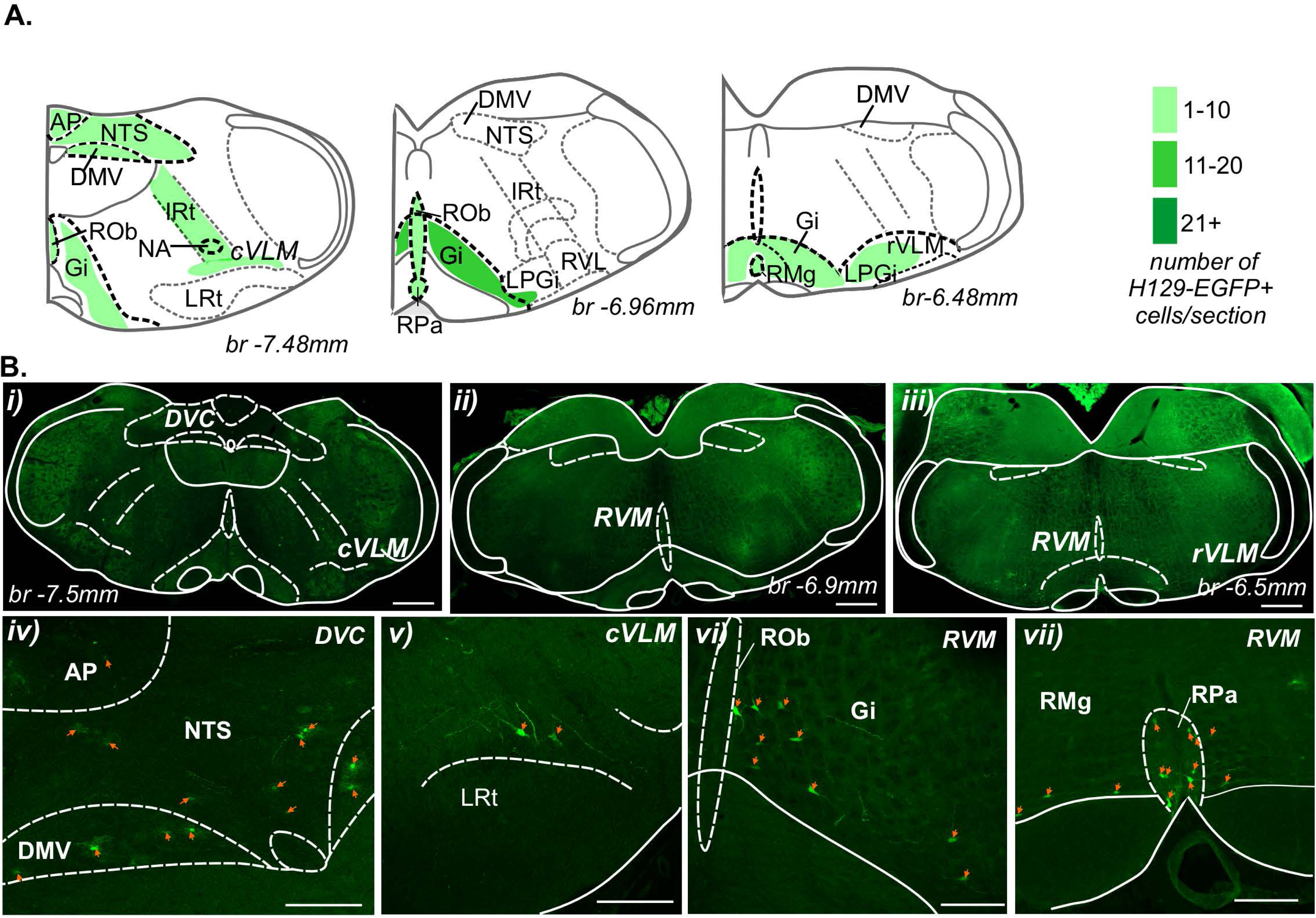
Distribution of H129-EGFP+ cells in the medulla 96 hours post-colorectal HSV-1 H129-EGFP inoculation. **A)** Schematic representation of the medulla (running caudal to rostral in the left to right axis) (Paxinos *et al*. 2001) summarizing distribution (regions with indicated by green shading) and semi-quantification of the density (indicated by intensity of green shading) of H129-EGFP+ cells 96-hours following HSV-1 H129-EGFP colorectal inoculation. Data obtained from 2-3 sections/ mouse, from N=3 female mice. **B)** Photomicrographs at **(i-iii)** low and **(iv-vii)** high magnification of **i)** caudal, **ii)** intermediate and **iii)** rostral sections of the medulla showing the distribution of H129-EGFP+ cells (green, highlighted by orange arrows) in the **(i, iv)** dorsal vagal complex (DVC) and within the reticulum of the **(i, v)** caudal ventrolateral medulla (cVLM) and **(ii, iii, vi, vii)** rostral ventromedial medulla (RVM) 96-hours post-inoculation of HSV-1 H129-EGFP within the colorectal wall. Scale bars: **i-iii:** 500 μm and **iv-vi:** 100 μm. Abbreviations: AP = Area postrema, br = bregma, DMV = dorsal motor nucleus of the vagus, Gi = gigantocellular reticular nucleus, IRt = intermediate reticular nucleus, NA = nucleus ambiguus, NTS = nucleus of the solitary tract, LRt = lateral reticular nucleus, LPGi = lateral paragigantocellular nucleus, rVLM = rostroventrolateral reticular nucleus, ROb= raphe obscurus nucleus, RMg = raphe magnus nucleus and RPa= raphe pallidus nucleus (RPa).

**Figure 5:**
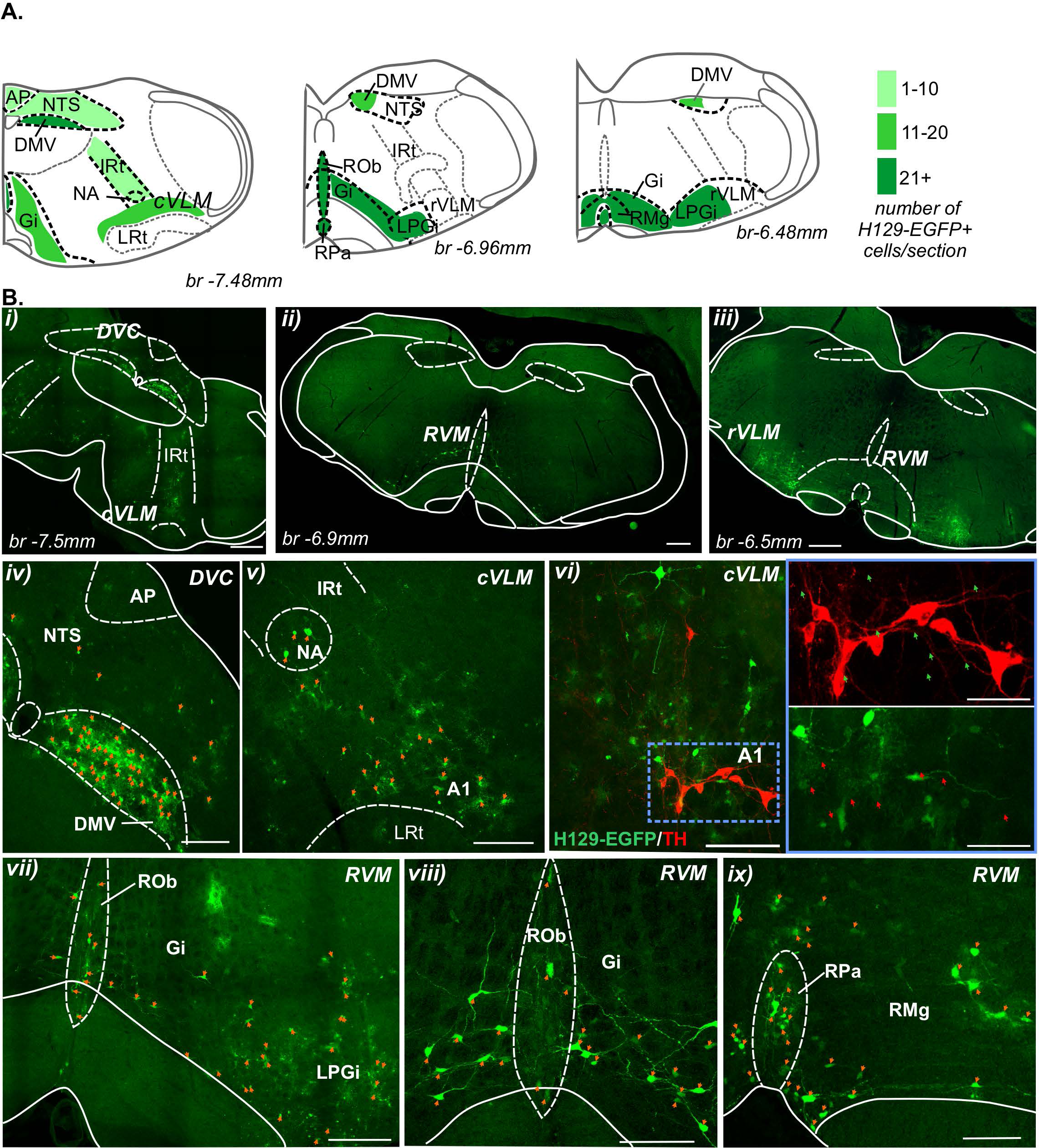
Distribution of H129-EGFP+ cells in the medulla 120 hours post-colorectal HSV-1 H129-EGFP inoculation. **A)** Schematic representation of the medulla oblongata (running caudal to rostral in the left to right axis) (Paxinos *et al*. 2001) summarizing distribution (regions with indicated by green shading) and semi-quantification of the density (indicated by intensity of green shading) of H129-EGFP+ cells 120-hours following HSV-1 H129-EGFP colorectal inoculation. Data obtained from 2-3 sections/mouse, from N= 8 mice (7F:1M) **B)** Photomicrographs at **(i-iii)** low and **(iv-x)** high magnification of **(i, iv, v, vi)** caudal, **(ii, vii, viii)** intermediate and **(iii, ix)** rostral sections of the medulla showing the distribution of H129-EGFP+ cells (green, highlighted by orange arrows) in the **(i, iv)** dorsal vagal complex (DVC), **(i, v, vi)** caudal ventrolateral medulla (cVLM) and **(ii, iii, vii, viii, ix)** rostral ventrolateral medulla (rVLM) and ventromedial medulla (RVM) 120-hours post-inoculation of the colorectal wall with HSV-1 H129-EGFP. **(vi)** Photomicrograph of the cVLM immunolabelled for tyrosine hydroxylase (TH, red), EGFP-labelled cells are indicated by green arrows and TH-immunoreactive neurons are indicated by red arrows. Images in the blue lined box (scale bar: 50 μm) correspond to the region within the blue dotted box. Scale bars: **i-iii:** 500 μm and **iv-ix:** 150 μm. Abbreviations: AP = Area postrema, br = bregma, DMV = dorsal motor nucleus of the vagus, Gi = gigantocellular reticular nucleus, IRt = intermediate reticular nucleus, NA = nucleus ambiguus, NTS = nucleus of the solitary tract, LPGi = lateral paragigantocellular nucleus, ROb= raphe obscurus nucleus, RMg = raphe magnus nucleus and RPa= raphe pallidus nucleus (RPa).

**Figure 6:**
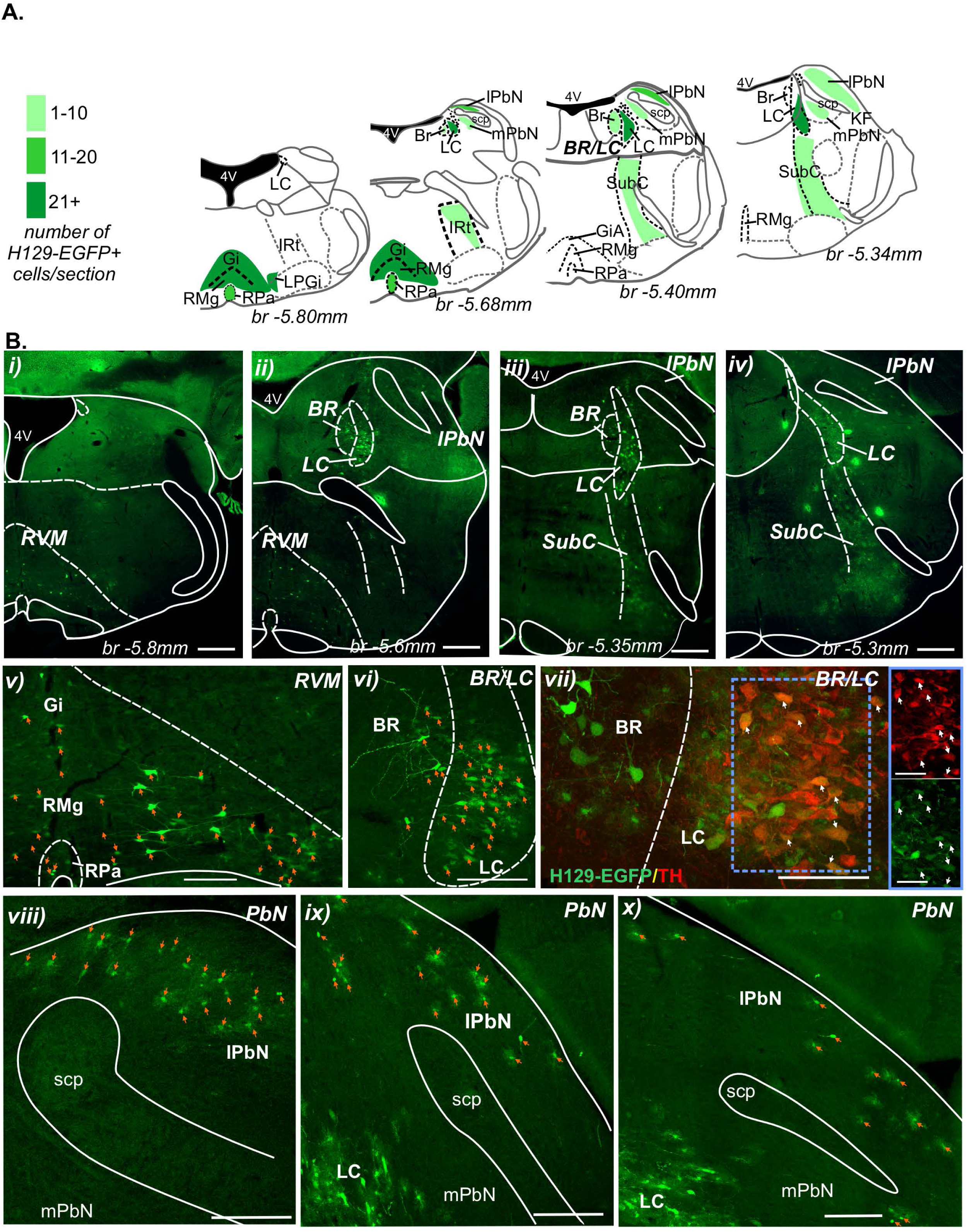
Distribution of H129-EGFP+ cells in the pons 120 hours post-colorectal HSV-1 H129-EGFP inoculation. **A)** Schematic representation of the pons (running caudal to rostral in left to right axis) (Paxinos *et al*. 2001) summarizing distribution (regions with indicated by green shading) and semi-quantification of the density (indicated by intensity of green shading) of H129-EGFP+ cells 120 hours following HSV-1 H129-EGFP colon inoculation. Gained from 2-3 sections/pons level/mouse, from N= 7 female mice. **B)** Photomicrographs at **(i-iv)** low and **(v-x)** high magnification of pontine sections from mice at 120 hours post-inoculation of HSV-1 H129-EGFP within the colorectal wall. Images demonstrate the distribution of H129-EGFP+ cells (green, highlighted by orange arrows) within the **(i, ii, v)** rostral ventromedial medulla (RVM), **(ii-iv, vi, vii)** Barrington’s nucleus (BR), locus coeruleus (LC) and sub coeruleus (subC) and the **(ii-iv, viii-x)** lateral parabrachial complex (lPbn). **(vii)** Photomicrograph of the BR and LC immunolabelled for tyrosine hydroxylase (TH, red), EGFP/TH co-labelled neurons are highlighted by white arrows. Image in the blue lined box (scale bar: 20 μm) corresponds to the region within the blue dotted box. Scale bars: **i-iv:** 500 μm, **v, vi, viii-x:** 150 μm and **vii:** 50 μm. Abbreviations: 4V = Fourth ventricle, br = bregma, Gi = gigantocellular reticular nucleus, IRt = intermediate reticular nucleus, KF = Kölliker-Fuse nucleus, lPbN = lateral parabrachial nuclei, LPgi = lateral paragigantocellular nucleus, mPbN = medial parabrachial nuclei, RMg = raphe magnus nucleus, RPa = raphe pallidus nucleus and scp = superior cerebellar peduncle.

**Figure 7:**
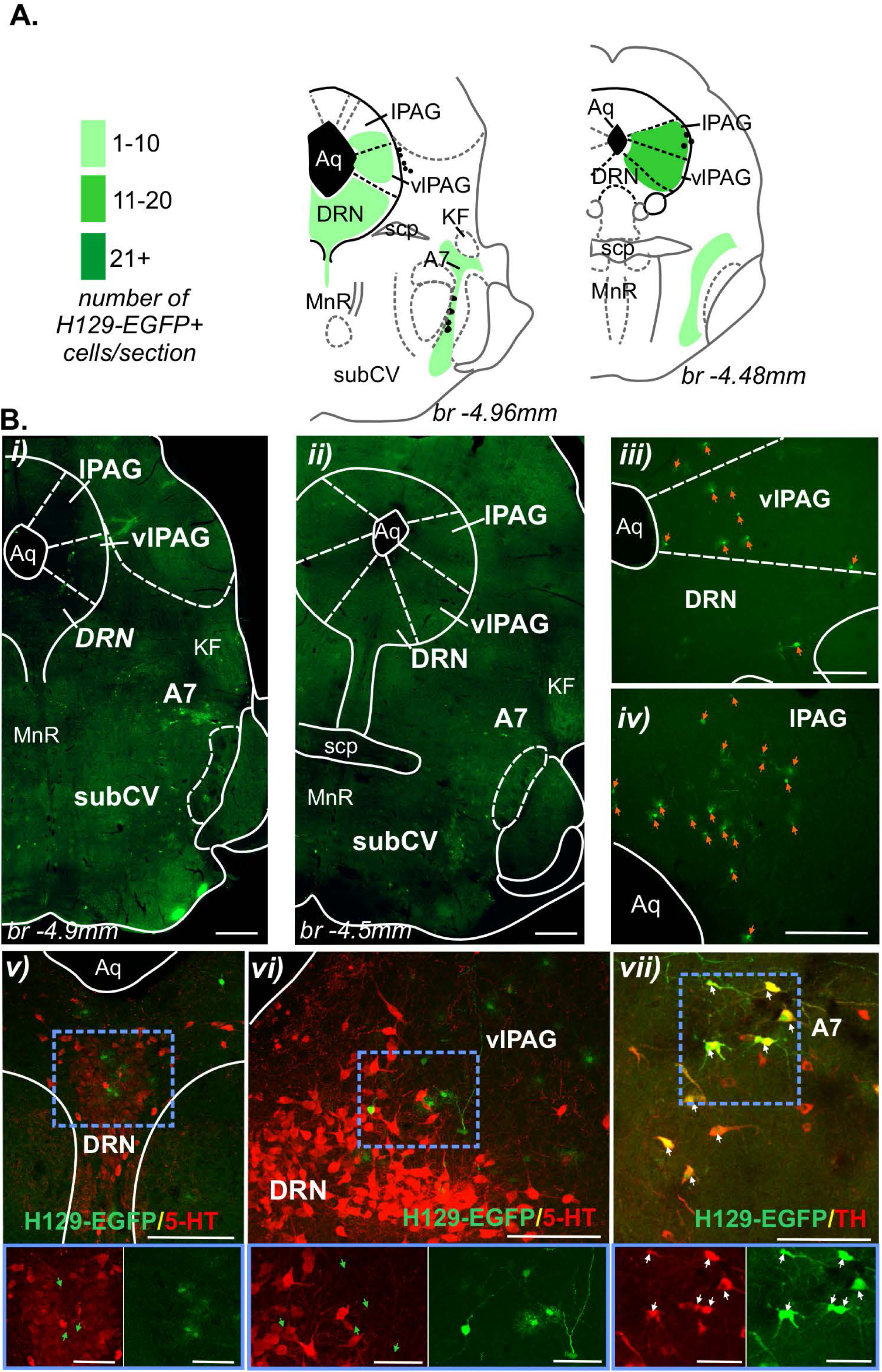
Distribution of H129-EGFP+ cells in the midbrain 120 hours post-colorectal HSV-1 H129-EGFP inoculation. **A)** Schematic representation of the caudal midbrain (running caudal to rostral in left to right axis) (Paxinos *et al*. 2001) summarizing distribution (regions with indicated by green shading) and semi-quantification of the density (indicated by intensity of green shading) of H129-EGFP+ cells 120 hours following HSV-1 H129-EGFP colon inoculation. Data obtained from 2-3 sections/midbrain level/mouse. 120 hours: N= 5 female mice. **B)** Photomicrographs at **(i-ii)** low and **(iii-vii)** high magnification of midbrain sections from mice at 120 hours post-inoculation of HSV-1 H129-EGFP within the colorectal wall, demonstrating the distribution of H129-EGFP+ cells (green, highlighted by orange arrows) within the **(i-vi)** periaqueductal gray (PAG) and the dorsal raphe nuclei (DRN), and the **(i, ii, vii)** A7. **(v, vi)** Photomicrographs of the PAG and DRN immunolabelled for serotonin (5-HT, red). H129-EGFP+ cells are indicated by green arrows. Image in the blue lined box (scale bar: 50 μm) corresponds to the region within the blue dotted box. **(vii)** Photomicrograph of the A7 immunolabelled for tyrosine hydroxylase (TH, red), EGFP/TH co-labelled neurons are highlighted by white arrows. Images in blue lined box (scale bar: 50 μm) correspond to the region within the blue dotted box. Scale bars: **i-ii:** 500 μm and **iii-vii:** 100 μm. Abbreviations: br = bregma, Aq = aqueduct, KF = Kölliker-Fuse nucleus, lPAG = lateral periaqueductal gray, MnR = median raphe nucleus, SubCV = sub-coeruleus ventral, scp = superior cerebellar peduncle and vlPAG = ventrolateral periaqueductal gray.

In the TL spinal cord, in tissue collected 72 hours post-inoculation H129-EGFP+ cells **(Fig. 2Ai)** were scattered within the superficial dorsal horn lamina I (LI, **Fig. 2Bi, 2Ci and 2Di)** and the intermediolateral nuclei (IML, **Fig. 2Bi, and 2Ci)**. In tissue collected 96-120 hours post-inoculation **(Fig. 2Aii and 2Aiii)**, H129-EGFP+ cells were additionally scattered in the deeper dorsal horn laminae III and V **(Fig. 2Bii-iii, 2Cii-iii, 2Dii-iii)**. Dense H129-EGFP+ labelling was evident in the ventrolateral edges of the ventral horn (vh) and within white matter tracts **(Fig. 2Aii and 2Div)**. In the LS spinal cord **(Fig. 3)**, H129-EGFP+ cells were scattered in the dorsal horn LI **(Fig. 3Ai, 3Bi, 3Ci and 3Di)** and the SPN **(Fig. 3Ai and 3Bi)** in tissue collected 72 hours post-inoculation. In tissue collected 96 hours post-inoculation **(Fig. 3Aii)**, dense H129-EGFP+ cells were distributed throughout LI-V **(Fig. 3Bii, 3Cii and 3Dii)**, the DGC **(Fig. 3Bii and 3Cii)** and the intercalated nuclei (ici, **Fig. 3Bii and 3Cii)**. In the LS spinal cord 120 hours post-inoculation **(Fig. 3Aiii)**, H129-EGFP+ cells were scattered throughout these locations **(Fig. 3Biii, 3Ciii and 3Diii)**, in addition to ventrolateral tracts of the ventral horn (vh) **(Fig. 3Biii and 3Ciii)**.

In the medulla collected 96 hours post-inoculation **(Fig. 4)**, H129-EGFP+ cells were scattered in the caudal medulla dorsal vagal complex (DVC, **Fig. 4A, 4Bi, 4Biv)**, in the reticulum of the caudal and rostral ventrolateral medulla (VLM, **Fig. 4A, 4Bi, 4Bv)** and the rostral ventromedial medulla (RVM) **(Fig. 4A, 4Bii-vii)**. In tissue collected 120 hours post-inoculation **(Fig. 5)**, H129-EGFP+ cells were denser within the caudal **(Fig. 5A, 5Bi)** and rostral medulla **(Fig. 5A, 5Bii, 5Biii)** relative to tissue collected at 96 hours. In the caudal DVC, H129-EGFP+ cells remained scattered in the area postrema (AP) and the nucleus of the solitary tract (NTS), yet were dense within the dorsal motor nucleus of the vagus (DMV) **(Fig. 5A, 5Bi, 5Biv).** Within the caudal VLM, H129-EGFP+ cells were evident in the lateral reticulum in the area of the nucleus ambiguus (NA, **Fig. 5Bv)**, intermediate reticular nucleus (IRt, **Fig. 5Bi)** and the reticular formation dorsal to the lateral reticular nucleus (LRt, **Fig. 5Bi, 5Bv, 5Bvi)** surrounding catecholamine (tyrosine hydroxylase (TH)-immunoreactive) A1 cell group **(Fig. 5Bvi)**. In the rostral medulla, substantial numbers of H129-EGFP+ cells were evident in the reticulum of the ventrolateral medulla **(Fig. 5A, 5Biii)** and the RVM raphe obscurus nucleus (ROb), gigantocellular reticular nucleus (Gi), lateral paragigantocellular nucleus (LPGi, **Fig. 5Bii, 5iii, 5vii, 5viii)**, the raphe magnus nucleus (RMg), and the raphe pallidus nucleus (RPa, **Fig. 5Biii and 5Bix)**.

In the pons **(Fig. 6)**, dense H129-EGFP+ cells were evident in tissue collected 120 hours post-inoculation in the RVM reticulum **(Fig. 6A, 6Bi, 6Bii, 6Bv)**, Barrington’s nucleus (BR, **Fig. 6A, 6Bii, 6Biii, 6Bvi, 6Bvii)**, locus coeruleus (LC, **Fig. 6A, 6Bii, 6Biii, 6Biv, 6Bvi, 6Bvii)** and the sub coeruleus (SubC, **Fig. 6A, 6Biii and 6Biv)**. H129-EGFP+ cells were also observed in the dorsal aspect of the lateral parabrachial nuclei (lPbn, **Fig. 6A, 6Bviii-x)**. Within the caudal midbrain **(Fig. 7)**, H129-EGFP+ cells were scattered within the lateral (lPAG) and ventrolateral (vlPAG) sub-regions of the periaqueductal gray (PAG, **Fig. 7A, 7Bi-iv, 7Bvi)**. EGFP-cells were also evident in the dorsal raphe nuclei (DRN, **Fig. 7A, 7Bi-iii and 7Bv)**, the ventral aspect of the SubC **(Fig. 7A, 7Bi-ii)** and within the A7 catecholamine cellular region, where they co-labelled for TH **(Fig. 7A, 7Bi-ii, 7Bvii)**.

### 4.2 Distribution of pERK-immunoreactive (IR) neurons in the brainstem following noxious colorectal distension (CRD)

Whilst care was taken to avoid leakage of the viral tracer, it is a possibility that some off target sites see virus and results in labelling in the brainstem not relevant to colorectal circuitry. To assess this limitation, the distribution of neurons in the brainstem activated by *in vivo* CRD was used to identify brainstem nuclei functionally relevant to colorectal sensory processing and compared to the distribution of H129-EGFP+ cells. Following noxious CRD, pERK-IR neurons were observed within the DMV and NTS **(Fig. 8A, 8B, 8C)** and cVLM **(Fig. 8A, 8B, 8D)** of the caudal medulla. Contrasting the distribution of H129-EGFP cells, pERK-IR neurons were relatively dense within the NTS **(Fig. 8A, 8B, 8C)** and populations of pERK-IR neurons within the cVLM immunolabelled for TH **(Fig. 8Diii)**. In the rostral medulla **(Fig. 8A, 8Bii, 8B iii)**, pERK-IR neurons were similarly distributed to H129-EGFP+ cells within the rVLM **(Fig. 8A, 8Bii, 8Biii)** and RVM **(Fig. 8A, 8Bi, 8Biii, 8E)**. In the DVC **(Fig. 8C)**, cVLM (**Fig. 8D)** and the RVM **(Fig. 8E)** compared to no CRD controls **(Fig. 8Ci, 8Di, 8Ei)**, noxious CRD **(Fig. 8Cii, 8Dii, 8Eii)** evoked significant increases in the number of pERK-IR neurons, which was equivalent in male and female mice **(Fig. 8Ciii, 8Div, 8Eiii)**. A significant difference between female and male mice was observed in the DMV, with a greater number of pERK-IR neurons evident in female mice following noxious CRD **(Fig. 8Ciii)**.

**Figure 8:**
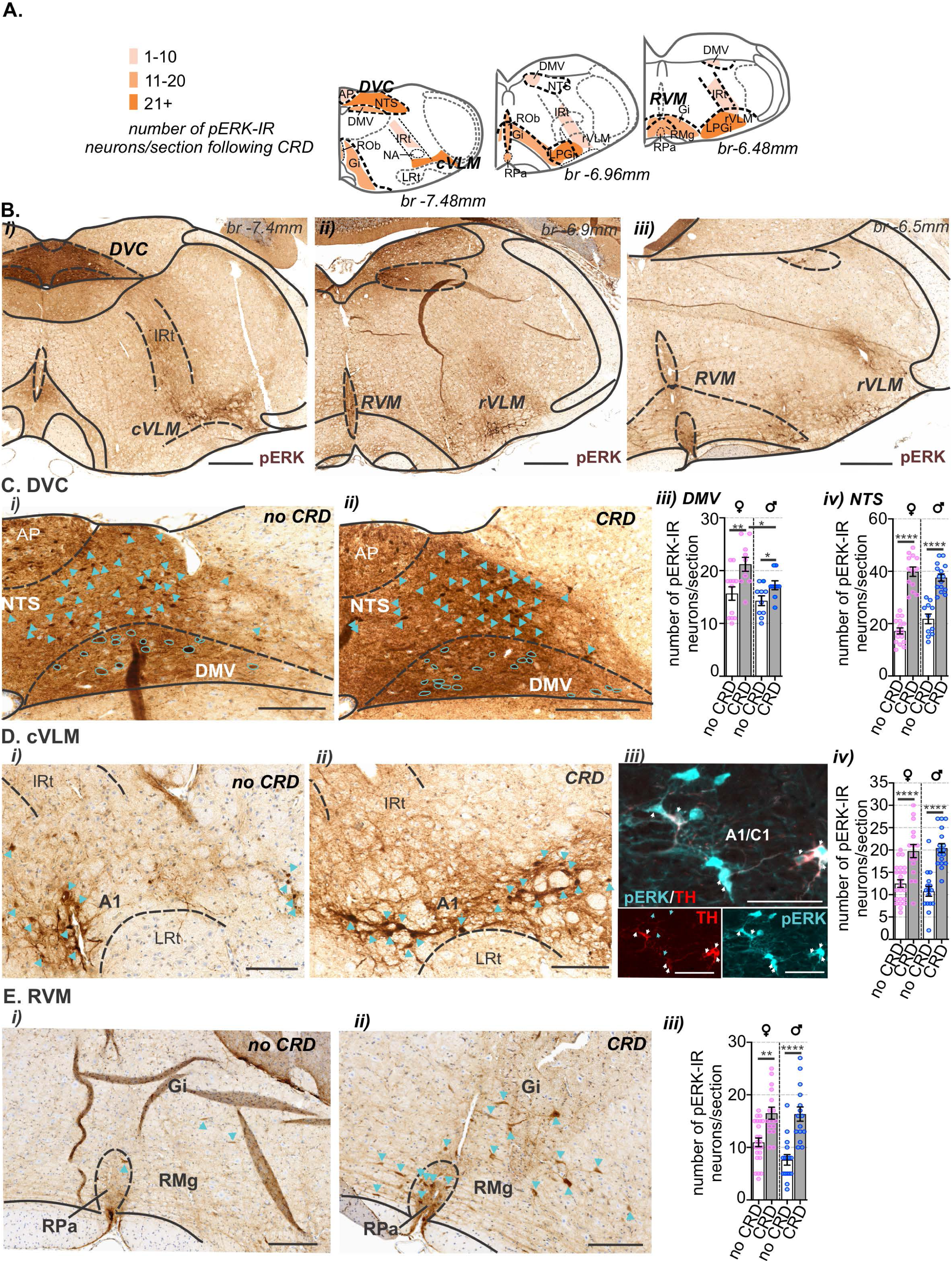
Distribution of pERK-immunoreactive (pERK-IR) neurons in the medulla following *in vivo* colorectal distension (CRD). **A)** Schematic representation of the medulla oblongata (Paxinos *et al*. 2001) summarizing the distribution (regions indicated by orange shading) and density (indicated by intensity of orange shading) of pERK-IR neurons following *in vivo* noxious CRD. Data obtained from N=9 mice (4F:5M). **B)** Low magnification photomicrographs of **(i)** caudal, **(ii)** intermediate and **(iii)** rostral sections of medulla showing the distribution of pERK-IR neurons (brown) following *in vivo* CRD at noxious pressures. Scale bars: 500 μm. **C-E)** High magnification photomicrographs showing the distribution of pERK-IR neurons (brown, indicated by blue outline or blue arrows) within the **C)** caudal dorsal vagal complex (DVC), **D)** caudal ventrolateral medulla (cVLM) and **E)** rostral ventromedial medulla (RVM) from mice that underwent **(i)** no CRD or **(ii)** noxious CRD (CRD). **(Diii)** Photomicrograph of the A1 nuclei within the cVLM immunolabelled for pERK (cyan) and tyrosine hydroxylase (TH, red), pERK-immunoreactive neurons are indicated by cyan arrows and pERK/TH co-labelled cells are indicated by white arrows. Scale bars: = 100 μm. Quantification comparing the number of pERK-IR neurons between female (♀, pink data points) and male (♂, blue data points) mice that underwent no CRD (white bar) or noxious CRD (grey bar) in sections of **C)** DVC, **(Ciii)** dorsal vagal motor nuclei (DMV) and **(Civ)** nucleus of the solitary tract (NTS), **(Diii)** cVLM (inclusive of the A1 region ventral to the intermediate reticular (IRt) and dorsal to the lateral reticular nucleus (LRt)) and **(Eiii)** RVM (inclusive of the raphe pallidus nucleus (RPa), raphe magnus nucleus (RMg) and the gigantocellular reticular nucleus (Gi)). * = *P*<0.05, ** = *P*<0.01, **** = *P*<0.0001, determined by One-Way ANOVA (parametric data) with Bonferroni multiple comparison tests. Individual data points represent the number of pERK-IR neurons in a section, gained from 2-5 sections/mouse from No CRD: N=10 mice (5F:5M) and CRD: N=9 mice (4F:5M). Abbreviations: AP = Area postrema, br = bregma, NA = nucleus ambiguus, LRt = lateral reticular nucleus, LPGi = lateral paragigantocellular nucleus, rRVM = rostroventrolateral reticular nucleus and ROb= raphe obscurus nucleus.

In the pons **(Fig. 9)**, and similar to H129-EGFP+ cells, pERK-IR neurons were evident within BR **(Fig. 9A, 9Bi-ii, 9C)**, the LC **(Fig. 9A, 9Bi-ii, 9C)**, the IRt and subC **(Fig. 9A, 9Bi-iii)** and the lPbN **(Fig. 9A, 9Bii-iii, 9D)**. In the BR **(Fig. 9C)**, relative to no CRD controls **(Fig. 9Ci)**, the number of pERK-IR neurons was greater following noxious CRD **(Fig. 9Cii).** This increase was significant in male mice but not female mice **(Fig. 9Civ)**. Conversely, in the LC **(Fig. 9C)**, relative to no CRD controls **(Fig. 9Ci)**, the number of pERK-IR neurons increased following noxious CRD **(Fig. 9Cii)** significantly within female mice but not male mice **(Fig. 9Cv)**. In the lPbN **(Fig. 9D)**, relative to no CRD controls **(Fig. 9Di)**, the number of pERK-IR neurons was significantly increased following noxious CRD **(Fig. 9Dii)**, which was equal in male and female mice **(Fig. 9Div)**.

**Figure 9:**
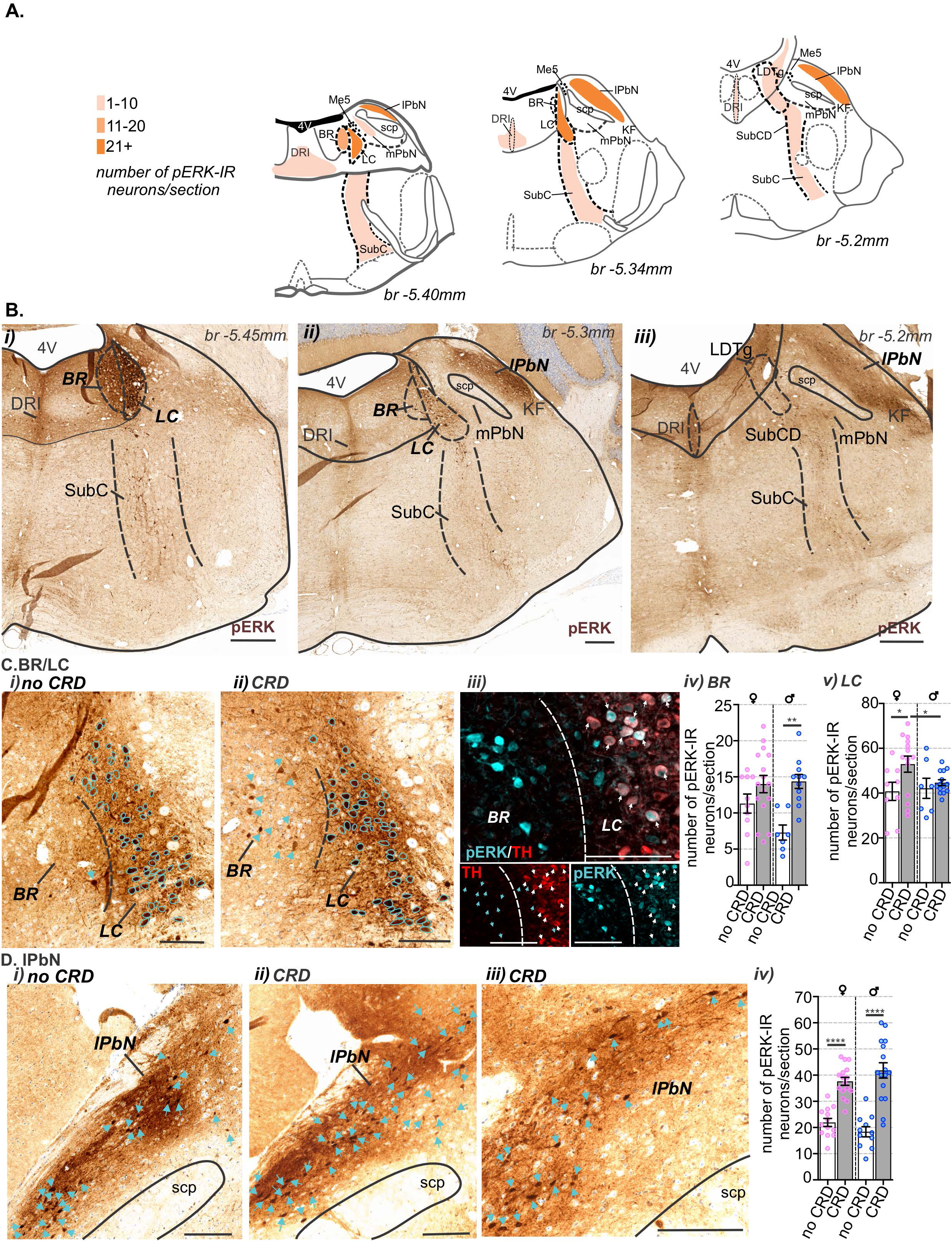
Distribution of pERK-immunoreactive (pERK-IR) neurons in the pons following *in vivo* colorectal distension (CRD). **A)** Schematic representation of the pons (Paxinos *et al*. 2001) summarizing the distribution (regions indicated by orange shading) and density (indicated by intensity of orange shading) of pERK-IR neurons following *in vivo* noxious CRD. Data obtained from N= 10 mice (5F:5M). **B)** Low magnification photomicrographs of **(i)** caudal, **(ii)** intermediate and **(iii)** rostral pontine sections showing the overall distribution of pERK-IR neurons (brown) following *in vivo* CRD at noxious pressures. pERK-IR neurons were present within the **(i, ii)** Barrington’s nucleus (BR), locus coeruleus (LC), intermediate reticular tracts (IRt), the **(ii, iii)** sub coeruleus (SubC), lateral parabrachial nucleus (lPbN) and **(i-iii)** in the dorsal raphe nuclei (interfascicular; DRI) and the **(iii)** laterodorsal tegmental nucleus (LDTg). Scale bars: 500 μm. **(C-D)** High magnification photomicrographs showing the distribution of pERK-IR neurons (brown, indicated by blue outline or blue arrows) within the **C)** BR and LC (BR/LC) and **D)** lPbN from mice that underwent **(i)** no CRD or **(ii-iii)** noxious CRD (CRD). **(Ciii)** Photomicrograph of the BR/LC immunolabelled for pERK (cyan) and tyrosine hydroxylase (TH, red), pERK/TH co-labelled cells are indicated by white arrows. Scale bars = 100 μm. Quantification comparing the number of pERK-IR neurons between female (♀, pink data points) and male (♂, blue data points) mice that underwent no CRD (white bar) or noxious CRD (grey bar)) within the **(Civ)** BR, **(Cv)** LC and the **(Dv)** lPbN. \**P*<0.05, **P<0.01, \*\*\*\**P*<0.0001, determined by One-Way ANOVA (parametric data) with Bonferroni multiple comparison tests. Individual data points represent the number of pERK-IR neurons in a section, gained from 2-4 sections/mouse from No CRD and CRD: N=10 mice (5F:5M). Abbreviations: 4V = Fourth ventricle, br = bregma, KF = Kölliker-Fuse nucleus, mPbN = medial parabrachial nuclei, Me5 = mesencephalic trigeminal nucleus, scp = superior cerebellar peduncle and SubCD = sub coeruleus dorsal.

In midbrain **(Fig. 10)**, aligning with the distribution of H129-EGFP+ cells, pERK-IR neurons were observed within the ventrolateral and lateral subregions of the PAG **(Fig. 10A, 10Bi-iii, 10Biv, 10C)**, the DRN **(Fig. 10A and 10Bi-iii**), the ventral SubC **(Fig. 10A and 10Bi),** and within the A7 **(Fig. 10A and 10Bi)**. Similar to H129-EGFP+ cells, pERK-IR neurons in the A7 co-labelled for TH **(Fig. 10Bv).** In the PAG compared to no CRD **(Fig. 10Ci)**, noxious CRD **(Fig. 10Cii)** evoked a significant increase in the number of pERK-IR neurons **(Fig. 10Ciii)**, which was equal in female and male mice.

**Figure 10:**
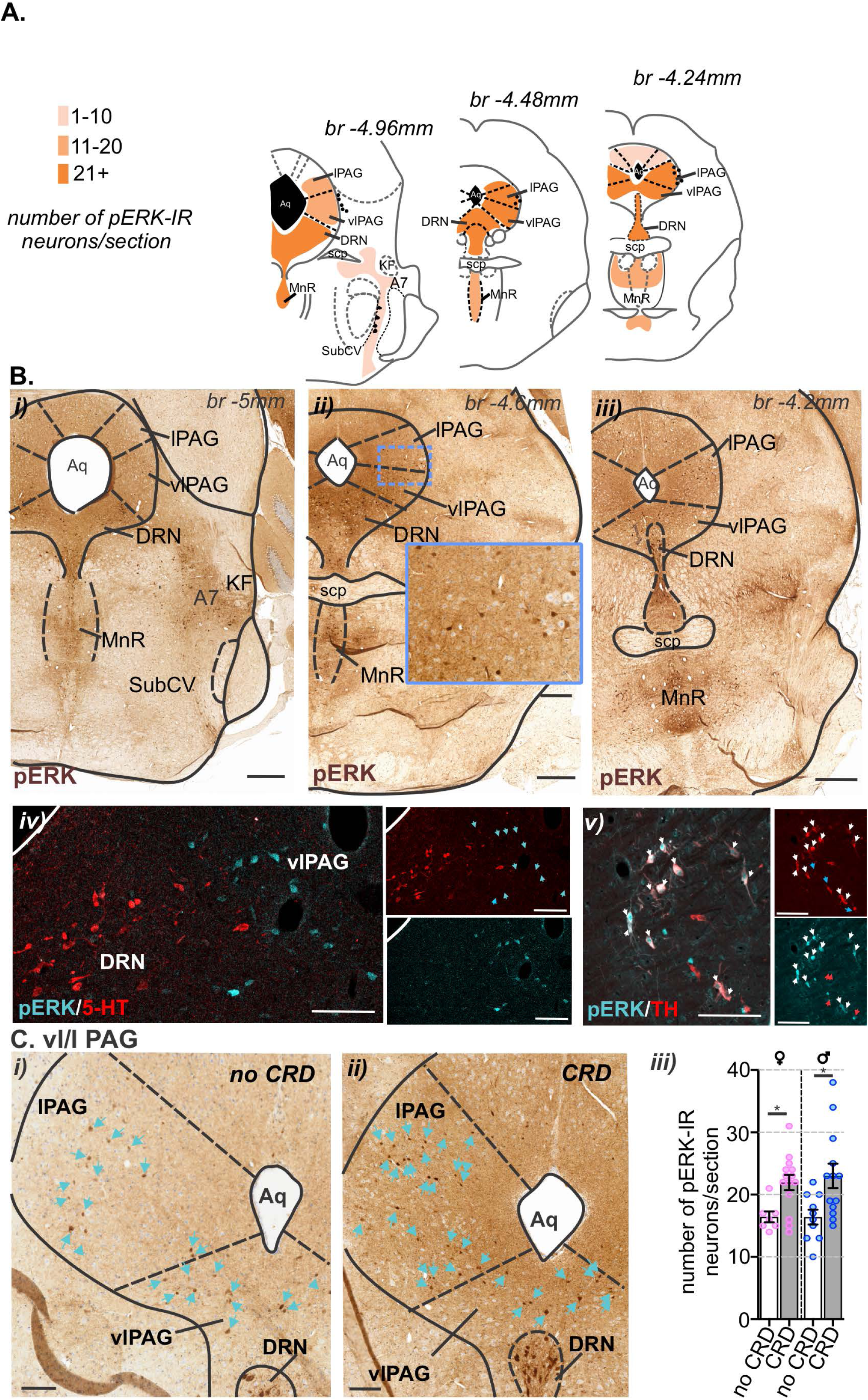
Distribution of pERK-immunoreactive (pERK-IR) neurons in the midbrain following *in vivo* colorectal distension (CRD). **A)** Schematic representation of the caudal midbrain (Paxinos *et al*. 2001) summarizing the distribution (regions indicated by orange shading) and density (indicated by intensity of orange shading) of pERK-IR neurons following *in vivo* noxious CRD. Data obtained from N= 10 mice (5F:5M). **Bi-iii)** Low magnification photomicrographs of caudal midbrain sections showing the overall distribution of pERK-IR neurons (brown) following *in vivo* CRD at noxious pressures. pERK-IR neurons were present within the **(i-iii)** periaqueductal gray (PAG), in the lateral (lPAG) and ventrolateral (vlPAG) sub-regions, in addition to the dorsal raphe nuclei (DRN) and the median raphe nucleus (MnR). **(ii)** Inset image (Scale bar = 100 μm) corresponds to region within dotted boxed area. Scale bars = 500 μm. **Biv)** High magnification photomicrographs of the vlPAG/DRN region immunolabelled for pERK (cyan) and serotonin (5-HT, red), pERK labelled cells are indicated by cyan arrows. Scale bars: 100 μm. **Bv)** High magnification photomicrographs of the A7 noradrenaline cellular region immunolabelled for pERK (cyan, cyan arrows) and tyrosine hydroxylase (TH, red, red arrows), pERK/TH co-labelled cells are indicated by white arrows. Scale bars: 100 μm. **C)** High magnification photomicrographs of the vl/lPAG region showing the distribution of pERK-IR neurons (brown, indicated by blue outline or blue arrows) within mice that underwent **(i)** no CRD or **(ii)** noxious CRD (CRD). Scale bars: 100 μm. **(iii)** Quantification comparing the number of pERK-IR neurons per section within the vl/lPAG between female (♀, pink data points) and male (♂, blue data points) mice that underwent no CRD (white bar) or noxious CRD (grey bar). \**P*<0.05 determined by One-Way ANOVA (parametric data) with Bonferroni multiple comparison tests. Individual data points represent the number of pERK-IR neurons in a section, gained from 2-3 sections/mouse from No CRD: N=9 mice (4F:5M) and CRD: N=10 mice (5F:5M). Abbreviations: Aq= aqueduct, br = bregma, KF = Kölliker-Fuse nucleus, Me5 = mesencephalic trigeminal nucleus, scp = superior cerebellar peduncle and SubCV = sub coeruleus ventral.

### 4.3 The effect of removing DRG at either TL or LS spinal levels on CRD evoked neuronal activation in the brainstem

Bilateral removal of the TL (T13-L1) or LS (L5-S1) DRG **(Fig. 11A)** relative to sham surgery (no DRG removal, **Fig. 11B)**, resulted in a reduction in the number of pERK-IR neurons within the dorsal horn at the respective levels of spinal cord **(Fig. 11C and 11D)**, as previously reported (Kyloh *et al*. 2022). In the DVC, relative to sham **(Fig. 12Ai)**, DRG removal at either spinal level, had no effect on the number of pERK-IR neurons evoked by CRD within the DMV, nor the NTS **(Fig. 12Aii-iv)**. Relative to the number of CRD evoked pERK-IR neurons in sham mice **(Fig. 12Bi, 12Ci, 13Ai, 13Bi and 13Ci)**, TL DRG significantly reduced pERK-IR neurons within the cVLM **(Fig. 12Bii and iv)**, the RVM **(Fig. 12Cii and iv)** and the lPbN **(Fig. 13Bii and iv)**, but not the BR and LC **(Fig. 13Aii and iv)** nor in the PAG **(Fig. 13Cii and iv)**. Whilst, LS DRG removal did not reduce the number of CRD evoked pERK-IR neurons in the cVLM **(Fig. 12Biii and iv)** but did significantly reduce pERK-IR number in the RVM **(Fig. 12Ciii and iv)**, the BR and the LC **(Fig. 13Aiii and iv)**, lPbN **(Fig. 13Biii and iv)** and the PAG **(Fig. 13Ciii and iv).**

**Figure 11:**
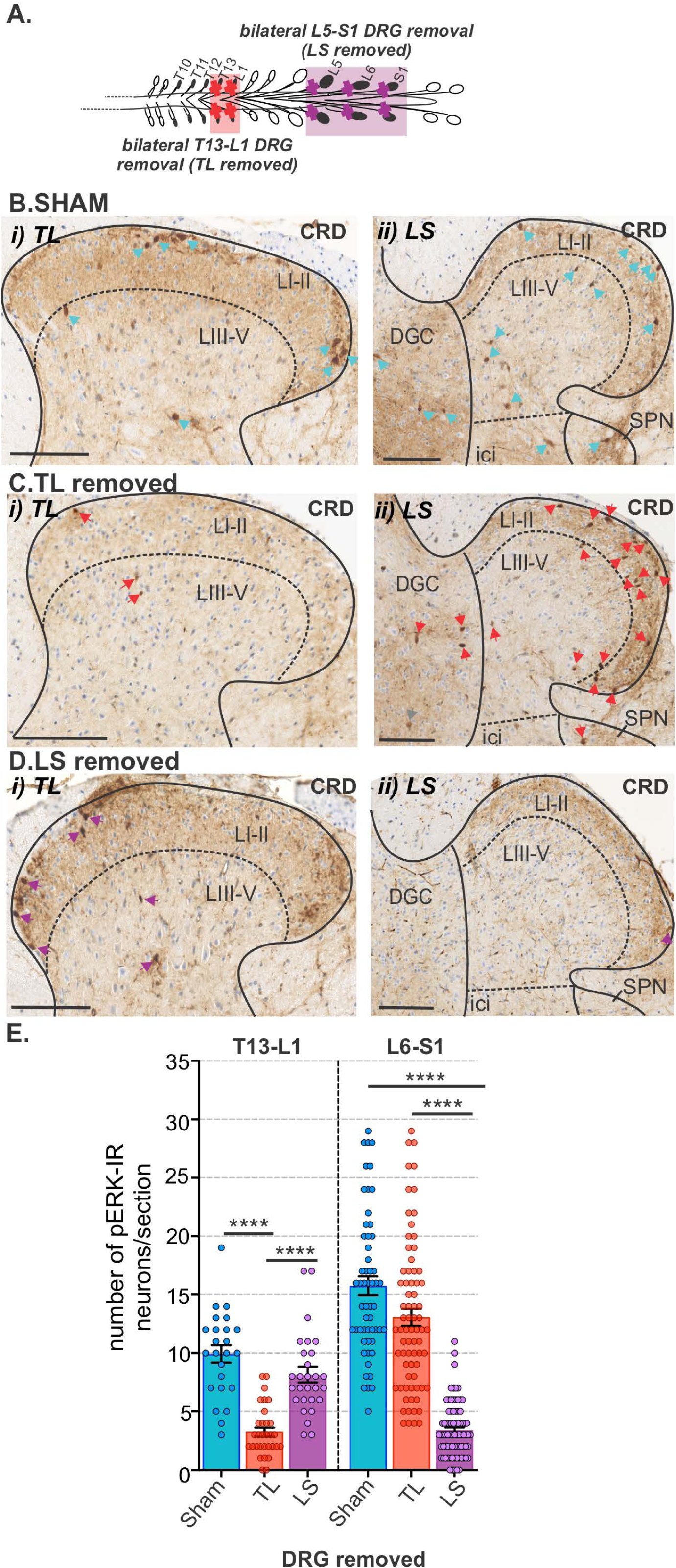
The effect of removing thoracolumbar (TL, T13-L1 DRG) and lumbosacral (LS, L5-S1) DRG on the number of pERK-IR neurons within the spinal cord following *in vivo* noxious colorectal distension. **A)** Schematic illustration showing the spinal levels at which dorsal root ganglia (DRG) were surgically removed in experimental groups TL removal (T13-L1 DRG, red box) and LS removal (L5-S1 DRG, purple box). The effect of either TL or LS DRG removal on neuronal activation (pERK-IR neurons) evoked by noxious CRD was compared to that in sham mice (that underwent the same surgical laminectomy procedure without DRG removal). **(B-D)** Photomicrographs showing the distribution of pERK-IR neurons (pERK, brown, indicated by arrows) within the **(i)** TL (T13-L1) and **(ii)** LS (L6-S1) spinal cord dorsal horn following noxious CRD in **B)** sham mice (blue arrows) and mice with **C)** TL (red arrows) or **D)** LS (purple arrows) DRG removed. Scale bars: 100 μm. **E)** Quantification comparing the number of pERK-IR neurons in the dorsal horn at TL (T13-L1) and LS (L6-S1) spinal levels between sham (blue), TL (red) and LS (purple) removed mice. \*\*\*\**P*<0.0001 determined by One-Way ANOVA (parametric data) with Bonferroni multiple comparison tests. Individual data points represent the number of pERK-IR neurons in a dorsal horn section (inclusive of LI-V, ici, DGC and IML/SPN), with 5 sections/mouse gained from N=5 female mice/group. Error bars represent the standard error of the mean. Abbreviations: LI= lamina I, LII= lamina 2, LIII= lamina 3, lamina IV= lamina 4, lamina V= lamina 5, ici= intercalated nuclei, IML= intermediolateral nuclei, DGC= dorsal grey commissure and SPN = sacral parasympathetic nuclei.

**Figure 12:**
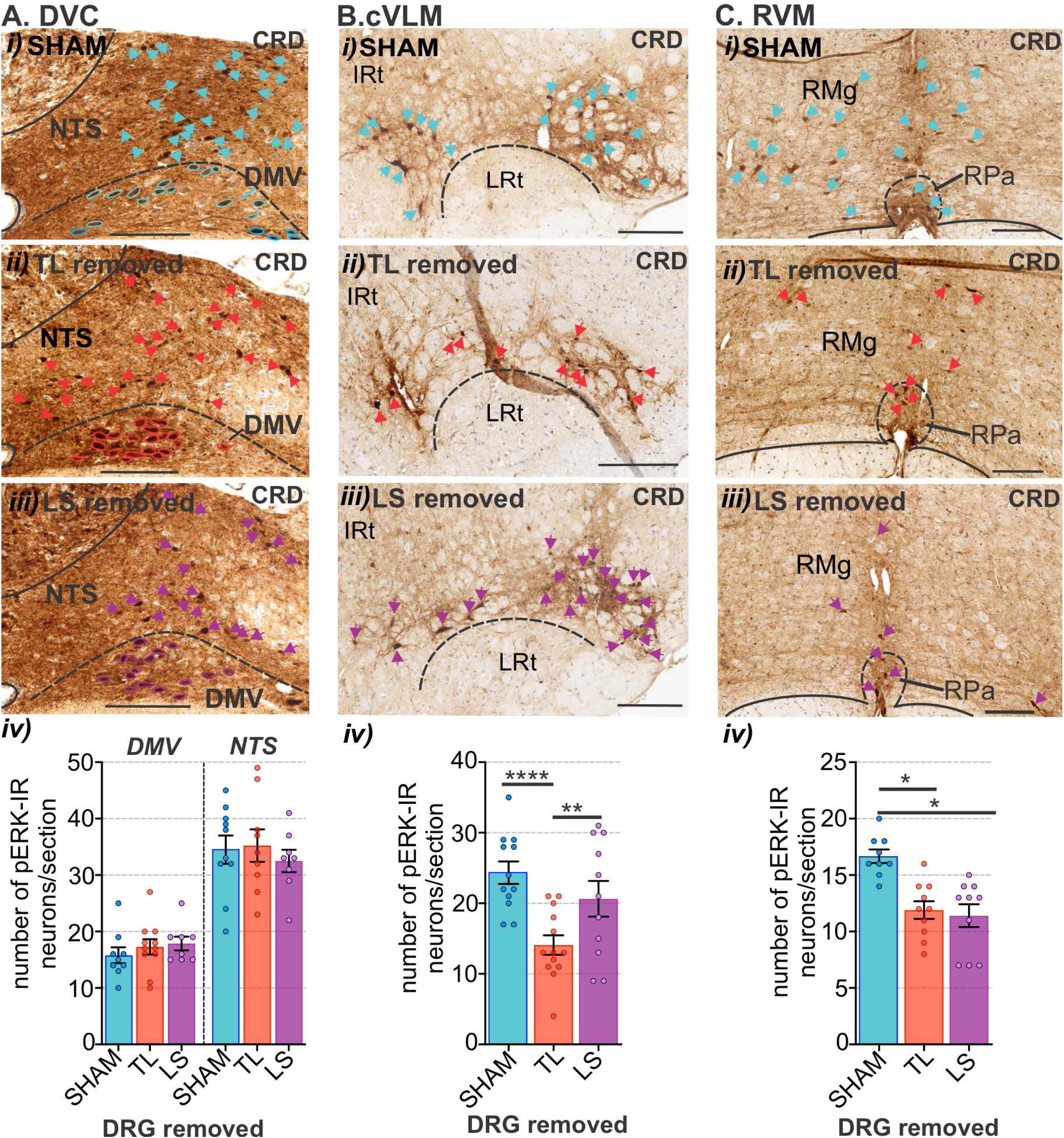
The effect of removing thoracolumbar (TL, T13-L1 DRG) and lumbosacral (LS, L5-S1) DRG on the number of pERK-IR neurons within the medulla following *in vivo* noxious colorectal distension. Photomicrographs showing pERK-IR neurons (pERK, brown, indicated by arrows) in the **A)** caudal dorsal vagal complex (DVC), **B)** caudal ventrolateral medulla (cVLM), and **C)** rostral ventromedial medulla (RVM) following noxious CRD in **(i)** sham mice (blue arrows) and mice with **(ii)** TL (red arrows) or **(iii)** LS (purple arrows) DRG removed. Scale bars: 100 μm. Quantification comparing the number of pERK-IR neurons between sham (blue), TL (red) and LS (purple) removed mice in the **(Aiv)** dorsal vagal motor nuclei (DMV) and nucleus of the solitary tract (NTS), **(Biv)** cVLM (inclusive of the A1 region ventral to the intermediate reticular (IRt) and dorsal to the lateral reticular nucleus (LRt)) and **(Civ)** RVM (inclusive of the raphe pallidus nucleus (RPa), raphe magnus nucleus (RMg)). \**P*<0.05, \*\**P*<0.01, \*\*\*\**P*<0.0001 determined by a One-way ANOVA with Bonferroni multiple comparison tests (parametric data). Individual data points represent the number of pERK-IR neurons per section from 1-2 sections/mouse from N=5 mice (female) for each of SHAM, TL removed and LS removed. Error bars represent the standard error of the mean.

**Figure 13:**
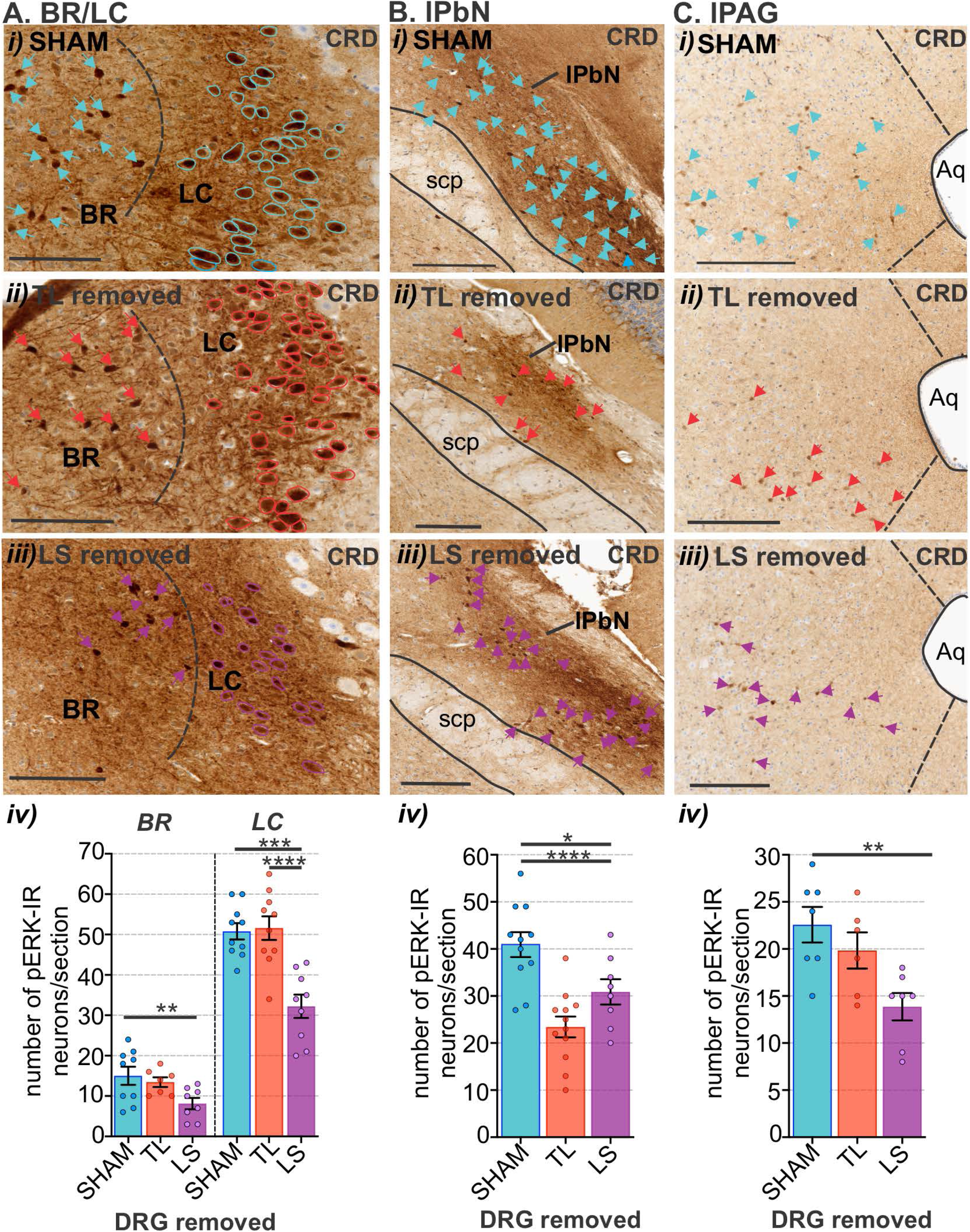
The effect of removing thoracolumbar (TL, T13-L1 DRG) and lumbosacral (LS, L5-S1) DRG on the number of pERK-IR neurons within the pons and midbrain following *in vivo* noxious colorectal distension. Photomicrographs showing pERK-IR neurons (pERK, brown, indicated by arrows) in the pontine **A)** Barrington’s nucleus (BR) and locus coeruleus (LC) and **B)** lateral parabrachial nucleus (lPbN), and the caudal midbrain **C)** periaqueductal gray (PAG), lateral (lPAG) sub-region, following noxious CRD in **(i)** sham mice (blue arrows) and mice with **(ii)** TL (red arrows) or **(iii)** LS (purple arrows) DRG removal. Scale bars: 100 μm. Quantification comparing the number of pERK-IR neurons in the **(Aiv)** Br and LC, **(Biv)** lPbN and **(Civ)** lPAG between sham (blue), TL (red) and LS (purple) removal mice. \**P*<0.05, \*\**P*<0.01, \*\*\**P*<0.001, \*\*\*\**P*<0.0001 determined by a One-way ANOVA with Bonferroni multiple comparison tests (parametric data). Individual data points represent the number of pERK-IR neurons per section from 1-2 sections/mouse from N=5 mice (female) for each of SHAM, TL removed and LS removed. Error bars represent the standard error of the mean. Abbreviations: Aq= aqueduct and scp = superior cerebellar peduncle.

## 5 Discussion

This study identified the brainstem structures relevant to colorectal sensory processing in the mouse and compared the relative contribution of ascending signalling via the TL and LS spinal cord on directing where colorectal sensory processing occurs within these structures. HSV1 H129-EGFP trans-neuronal tracing from the colorectum identified the spinal pathways and brainstem structures synaptically connected to the colorectum. H129-EGFP+ cells were observed in the dorsal horn of the TL and LS spinal cord 72 hours after colonic inoculation, in the medulla 96 hours after inoculation, and in pontine and midbrain nuclei 120 hours after inoculation **(Fig. 14A)**. Confirming their functional relevance to colorectal sensory processing, noxious CRD evoked increased neuronal activation, relative to no CRD, in many of these H129-EGFP+ brainstem structures **(Fig. 14B)**. Removal of TL DRG (splanchnic ascending relay) reduced neuronal activation in the cVLM, the RVM and the lPbN **(Fig. 14C)**. In comparison removal of LS DRG (pelvic ascending relay) resulted in a reduction of neuronal activation within the RVM, LC, BR, lPbN and the PAG **(Fig. 14D)**. Collectively, this data provides new insight into how the dual spinal afferent innervation of the colorectum differentially shapes where colorectal sensory information is integrated within brainstem circuits relevant to sensory and motor aspects of colorectal functions.

**Figure 14:**
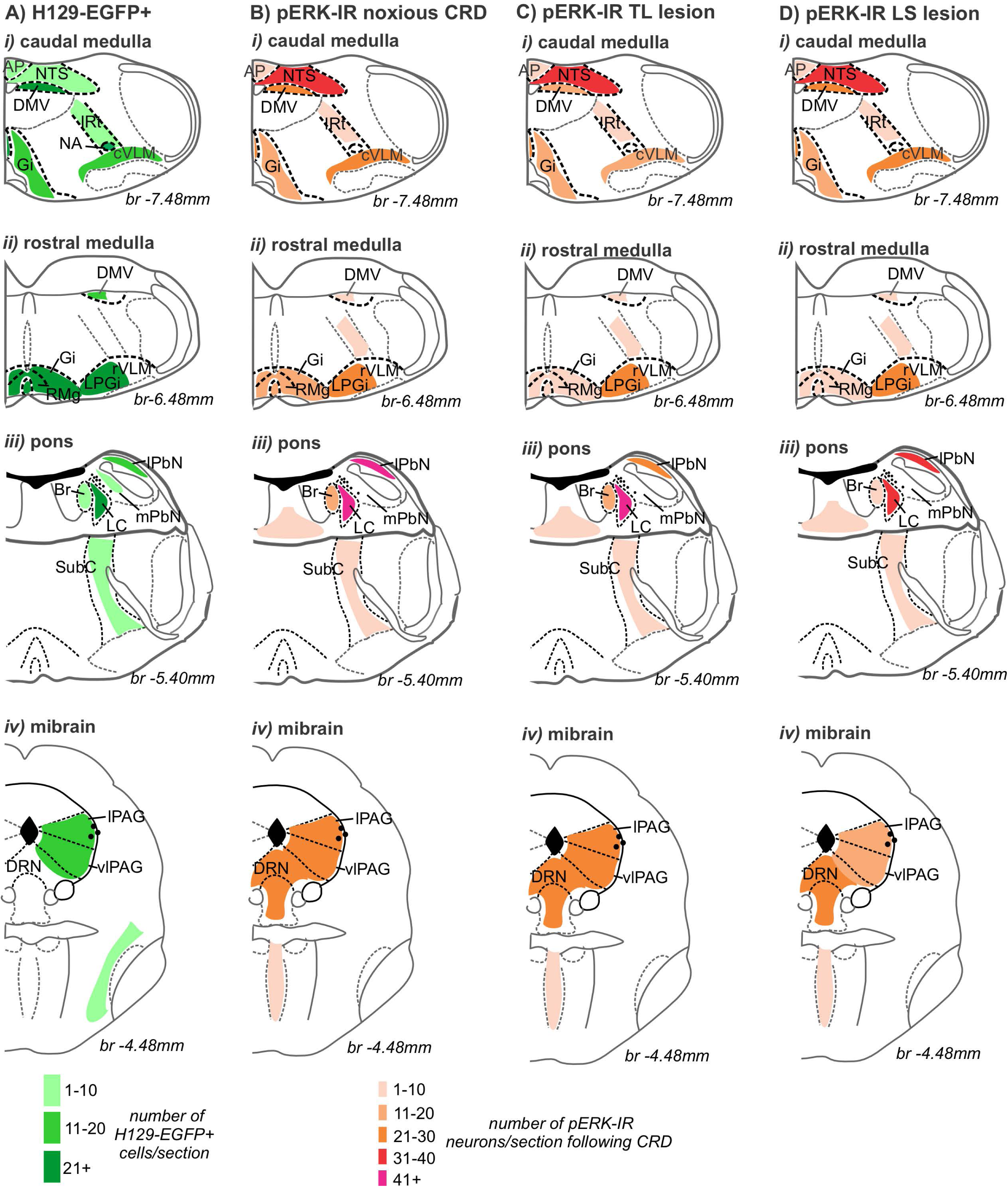
Summary diagram. Schematic (Paxinos *et al*. 2001) comparing of the distribution (shaded regions) and density (indicated by intensity of shading) of: **A)** H129-EGFP+ in tissue collected 120 hours after colorectal inoculation, **B)** pERK-IR neurons following *in vivo* colorectal distension (CRD) at noxious pressures. **C)** pERK-IR neurons following *in vivo* noxious CRD in mice that had TL DRG (T13-L1) removed and **D)** pERK-IR neurons following *in vivo* noxious CRD in mice that had LS DRG (L5-S1) removed in the ***i)*** caudal midbrain, ***ii)*** rostral medulla, ***iii)*** pons and ***iv)*** midbrain.

Within the TL and LS spinal cord dorsal horn, the distribution of H129-EGFP+ cells was consistent with that of colorectal afferent projections we have shown previously (Harrington *et al*. 2019; Wang *et al*. 2023). The exception being the abundance of labelling in the ventrolateral tracts in the TL spinal cord. We have not observed colorectal afferent projections nor pERK-IR neurons in this location in previous studies. It is a possibility that this labelling is a product of off target sites exposed to virus, or a consequence of retrograde tracing autonomic efferents. In the brainstem, H129-EGFP+ cells were evident within structures that are known to receive abundant input from the spinal cord, namely reticular formations within the caudal and rostral VLM and RVM, the lPbN, and the PAG (Lima *et al*. 1991; Wang *et al*. 1999; Polgar *et al*. 2010; Todd 2010; Martins *et al*. 2017; Wercberger *et al*. 2019; Gu *et al*. 2023). Whilst other labelled structures, such as the BR, LC and RVM, have been shown to receive direct input from the spinal cord, yet to a lesser degree in comparison to the aforemention regions (Cedarbaum *et al*. 1978; Ding *et al*. 1997; Wang *et al*. 1999; Samuels *et al*. 2008; Peng *et al*. 2023). Many of these structures also have descending outputs to the spinal cord which may have contributed to H129-EGFP+ labelling via retrograde transport (Samuels *et al*. 2008; Kawatani *et al*. 2021; Gu *et al*. 2023; Peng *et al*. 2023). The distribution pattern of H129-EGFP+ cells we observed in the brainstem aligned with findings from studies using pseudorabies virus (PRV) transneuronal tracing from the distal colon of the rat (Pavcovich *et al*. 1998; Valentino *et al*. 2000; Vizzard *et al*. 2000; Rouzade-Dominguez *et al*. 2003; He *et al*. 2018). However, unlike PRV, HSV-1 H129 has been shown to have a preferential, but not exclusive, anterograde direction of transport through central circuits (Rinaman *et al*. 2004; McGovern *et al*. 2012; McGovern *et al*. 2012; Wojaczynski *et al*. 2015; Parker *et al*. 2020). A study directly comparing HSV-1 H129 and PRV transport from the stomach wall illustrates that both PRV and HSV-1 produced first-order retrograde infection of sensory neurons and central autonomic neurons that project directly to peripheral inoculation sites (Rinaman *et al*. 2004). However, unlike PRV, subsequent HSV-1 transport through the spinal cord and brain was described as occuring predomominantly in the anterograde (ascending) direction via their synaptic output. As oppposed to transport in the retrograde direction from infected neurons into central sources via their synaptic inputs which was found to be limited (Rinaman *et al*. 2004). This distinction has particular relevance to our study, as we wanted to identify brainstem structures targeted by ascending pathways and limit the potential for retrograde labelling of brainstem structures via their descending projections onto HSV-1 H129 infected spinal cord neurons. While in our study this mode of transport cannot be completely excluded, a discrepancy between H129-EGFP+ and pERK-IR labelling of TH-IR catecholamine A1 neurons within the cVLM may support a preferential anterograde transport of HSV-1 H129. Populations of A1 catecholaminergic neurons activated by nociceptive stimuli have descending outputs in the spinal cord, but do not receive direct input from the spinal cord (Tavares *et al*. 2002; Gu *et al*. 2023). We observed many H129-EGFP+ labelled cells in the cVLM; however, these were not TH-IR catecholamine neurons. This is despite observing populations of pERK-IR within the cVLM that were A1 catecholaminergic neurons.

In the spinal cord dorsal horn, projection neurons primarily targeted by nociceptive afferent input that conveys into the brainstem are most dense within LI (Wercberger *et al*. 2019; Peirs *et al*. 2020). The primary brainstem targets of these neurons are reticular formations within the cVLM, the NTS, the lPbN and the PAG (Todd 2010; Cameron *et al*. 2015; Martins *et al*. 2017; Wercberger *et al*. 2019). Populations of dorsal horn neurons relevant to transmitting nociceptive and wide dynamic information are scattered throughout laminae III-VI and VII–VIII, and the DGC (Wercberger *et al*. 2019). These neurons terminate widely within the brainstem, in the NTS, in medullary and pontine reticular formation, the lPbN and the PAG (Todd 2010; Martins *et al*. 2017; Wercberger *et al*. 2019). Neurons within the SPN of the sacral spinal cord have shown to have ascending output to the BR (Ding *et al*. 1997; Wang *et al*. 1999), whilst there is evidence for preganglionic neurons within the IML of the thoracic spinal cord projecting into the LC (Cedarbaum *et al*. 1978). Based on this knowledge, in the TL spinal cord, H129-EGFP+ cells were distributed in laminae LI and LIII-V, which corresponds with our previous findings of a large proportion of splanchnic colorectal afferents being nociceptive and that they relay into dorsal horn circuits primary concerned with its supraspinal relay (Harrington *et al*. 2019; Wang *et al*. 2024). Illustrated by removal of TL DRG, this supraspinal output relays into the reticulum of the cVLM, and possibly the RVM, and the lPbN. Conversely, in the LS spinal cord dorsal horn H129-EGFP+ cells were more widely distributed within laminae LI-LV, DGC and SPN **(Fig. 14A)**. This reflects pelvic colorectal afferent fibres relaying a mix of sensory information (nociceptive and a wide dynamic range) into dorsal horn circuits underlying supraspinal relay, interneuronal processing and parasympathetic motor output, as we have described previously (Harrington *et al*. 2019; Wang *et al*. 2024). This is also reflected in the brainstem, with the effect of LS DRG removal indicating the information from colorectal pelvic afferents is relayed into multiple brainstem structures (RVM, lPbN, BR, LC and the PAG) that are involved in a diverse range of sensory and motor functions (Liu *et al*. 2007; Naitou *et al*. 2018; Nakamori *et al*. 2018; Lyubashina *et al*. 2022; Qi *et al*. 2022)

Comparing the effect of TL or LS DRG removal on colorectal processing within the brainstem, removal of either TL or LS DRG significantly reduced CRD-evoked neuronal activation within the lPbN and the RVM (Fig. 14C and D). As a major relay nucleus, the lPbN distributes nociceptive information widely within cortico-limbic and brainstem structures that shape pain behaviour, modulation and autonomic integration (Roeder *et al*. 2016; Chiang *et al*. 2020). The lPbN receives input from the spinal cord neurons distributed within the dorsal horn LI, and LIII-IV, which differentially contribute to nociceptive behaviour and autonomic responses (Choi *et al*. 2020; Browne *et al*. 2021). The effect we observed of TL DRG removal on the lPbN aligns with our recent findings of colorectal splanchnic afferent fibres in apposition to lPbN-projecting neurons within LI and that a large proportion (63%) of neurons activated by noxious CRD within LI of the TL spinal cord project to the lPbN (Wang *et al*. 2024). In the LS spinal cord, whilst lPbN-projection neurons make up a relative small proportion of the CRD responsive neurons in the dorsal horn LI (13%), lPbN-projecting neurons responsive to CRD have also been observed in the SPN, DGC and lateral grey matter (Murphy *et al*. 2009; Nishida *et al*. 2022; Wang *et al*. 2024). These differences indicates that the lPbN receives different qualities of colorectal information from the TL spinal cord as compared with the LS spinal cord. There is evidence of this, from the LS spinal cord at least, in an optogenetic study showing that populations of lPbN-projecting neurons within the DGC of the sacral spinal cord are involved in colitis evoked aversion responses but not visceromotor responses (Qi *et al*. 2022). The neurocircuits of the lPbN receiving differential colorectal input from the TL and LS spinal cord warrant further investigation, in particular assessing if they relay into disparate cortico-limbic circuits, that were not the focus of this study, to influence pain behaviours.

TL or LS DRG removal both reduced the number of RVM neurons activated by noxious CRD equally **(Fig. 14C and D)**. In the case of the TL pathway, this may be mediated via the lPbN as the RVM primarily receives ascending nociceptive information via the lPbN (Peng *et al*. 2023). Under physiological conditions, and when exposed to acute pain stimuli, the lPbN input into the RVM triggers acute hyperalgesia (Peng *et al*. 2023). As for the LS pathway, this may not solely be linked to reduced output from the lPbN, but also from the PAG and the LC, the activity of which was also reduced by LS DRG removal. Projections from PAG to RVM are involved in generating defensive responses to pain, whilst those from the LC are linked to both pro- and anti-nociception (Peng *et al*. 2023). Stimulation of the PAG inhibits CRD-evoked activity within LS spinal cord dorsal horn LI (Okada *et al*. 1999). This is mediated by the PAG relaying nociceptive input into the RVM and the LC with descending projections within LS spinal cord (Liu *et al*. 2008; Lyubashina *et al*. 2022; Lubejko *et al*. 2024). The PAG output to the LC and BR is also involved in recruiting neurons with descending output to the LS spinal cord defecation centre and the SPN to facilitate motility reflexes (Ding *et al*. 1997; Pavcovich *et al*. 1998; Nakamori *et al*. 2019). The ventrolateral and lateral subregions of the PAG, in which we observed H129-EGFP+ cells and neuronal activated by CRD, receive direct input from neurons in the sacral spinal cord distributed within LI, the lateral spinal nucleus and LV (Keay *et al*. 1997). The BR and LC also receive direct input from neurons distributed within the sacral spinal cord SPN and DGC, however to a lesser degree than that to the PAG (Ding *et al*. 1997; Wang *et al*. 1999). Our findings do provide clear evidence for the predominate role of pelvic afferent relay through the LS spinal cord to shaping colorectal sensory relay into and processing within the PAG, BR and LC. To clearly ascertain how much of this processing is shaped by the direct input from the spinal cord, the proportions of BR, LC or PAG projecting neurons in the LS spinal cord activated by colonic stimulation, and visa verse, is required. Interestingly, we observed a sex difference in the LC, with noxious CRD evoking an increase in pERK-IR neurons relative to no CRD controls in female mice but not male mice. This difference was not evident in the BR nor the PAG. The LC is known to exhibit sex differences in activation, which are mediated by sensitivity to corticotropin-releasing factor (Bangasser *et al*. 2016). This is thought to contribute to the greater vulnerability of females to stress-related disorders as the LC is involved in regulating levels of arousal by releasing norepinephrine into the forebrain (Bangasser *et al*. 2016). Our findings identify these sex differences in the LC may also affect colorectal nociception control and how it is linked with affective defecation reflexes.

DRG removal did not have an affect on CRD evoked neuronal activation in the NTS or in the DMV **(Fig. 14C and D)**. This provides insight into the influence the vagal pathways have on colorectal sensory processing within the NTS over that of spinosolitary tracts. In agreement, vagal denervation, rather than spinalization, has previously shown to significantly attenuate neuronal activation within the NTS, in addition to the rVLM, evoked from the proximal colon (Monnikes *et al*. 2003). We observed very few EGFP-H129+ cells within the NTS 96-120 hours after colorectal injection. The length of the balloon used for CRD would activate afferent fibres within the proximal and distal colon, whilst the HSV1-H129-EGFP injections were localised to the distal colon. The vagal afferent innervation of the colorectum occurs in a gradient. Anterograde tracing from the nodose ganglia in rats and mice show vagal afferent nerve endings within the proximal colon, that become sparse or are absent in the distal colon through to the rectum (Wang *et al*. 2000; Spencer *et al*. 2024). Aligning with this, a greater proportion of neurons within the nodose ganglia are retrogradely labelled from the proximal colon compared to the distal colon (Osman *et al*. 2023; Wang *et al*. 2024). In agreement, we have demonstrated that fibres in the NTS retrogradely labelled from the proximal colon are more abundant than fibres labelled from the distal colon (Wang *et al*. 2024). Moreover, fibres labelled from the proximal colon arborise extensively within the NTS, and are also present within the DMV, which may be reflected in the abundance of CRD evoked pERK-IR neurons in these locations. We have found that a proportion (13%) of CRD activated neurons in the NTS project to the lPbN (Wang *et al*. 2024). Based on the findings from this study, this is a minor contribution to the overall CRD evoked neuronal activation in the lPbN relative to spinal input. Unlike the NTS, we observed EGFP-H129+ cells in abundance within the DMV **(Fig. 14)**. This is possibly a consequence of retrograde labelling from DMV efferent projections that have been shown within the myenteric ganglia extending into the distal colon (Berthoud *et al*. 1991; Tao *et al*. 2021). Aligning with this, we have shown that DMV neurons are retrogradely labelled from the proximal and distal colon (Wang *et al*. 2024).

Collectively, this study identifies the brainstem structures involved in processing colorectal sensory information in the mouse. This study also highlights how brainstem structures are recruited by colorectal sensory processing and the similarities and differences induced by the splanchnic or pelvic spinal afferent pathways. These findings provide insight into the central circuitry to which the dual spinal afferent innervation of the colorectum is connected and aids interpretation of their role in affecting visceromotor responses and defecation associated with colorectal nociception. These findings also have implications for understanding how peripheral hypersensitivity or spinal sensitization within these spinal ascending pathways may lead to altered brainstem processing supporting hyperalgesia and autonomic dysfunction.

## 8 Additional information

### 8.2 Author Contributions

A.M.H, G.R, S.B.M, N.S and S.M.B devised experiments. A.M.H, QQ.W, A.E.M and M.K performed experiments and analyzed the data. A.M.H prepared figures and wrote the manuscript. G.R, S.M.B, S.B.M, A.E.M and N.S provided scientific input and corrections to the paper.

## 8.4 Acknowledgements

The authors acknowledge the instruments and expertise of Microscopy Australia (ROR: 042mm0k03) at Adelaide Microscopy, University of Adelaide, enabled by NCRIS, university, and South Australia government support. Parts of this work have appeared in presentations at the Australasian Neuroscience Society annual meeting 2019 and Society for Neuroscience: Global Connectome Virtual Conference 2021.

## 6 References

Asmus, S. E., E. K. Anderson, M. W. Ball, B. A. Barnes, A. M. Bohnen, A. M. Brown, . . . A. E. Warner (2008). Neurochemical characterization of tyrosine hydroxylase-immunoreactive interneurons in the developing rat cerebral cortex. Brain Res 1222: 95–105.

Bangasser, D. A., K. R. Wiersielis and S. Khantsis (2016). Sex differences in the locus coeruleus-norepinephrine system and its regulation by stress. Brain Res 1641(Pt B): 177–188.

Bankhead, P., M. B. Loughrey, J. A. Fernandez, Y. Dombrowski, D. G. McArt, P. D. Dunne, . . . P. W. Hamilton (2017). QuPath: Open source software for digital pathology image analysis. Sci Rep 7(1): 16878.

Barnett, E. M., G. D. Evans, N. Sun, S. Perlman and M. D. Cassell (1995). Anterograde tracing of trigeminal afferent pathways from the murine tooth pulp to cortex using herpes simplex virus type 1. J Neurosci 15(4): 2972–2984.

Bassi, J. K., A. A. Connelly, A. G. Butler, Y. Liu, A. Ghanbari, D. G. S. Farmer, . . . A. M. Allen (2022). Analysis of the distribution of vagal afferent projections from different peripheral organs to the nucleus of the solitary tract in rats. J Comp Neurol 530(17): 3072–3103.

Berthoud, H. R., N. R. Carlson and T. L. Powley (1991). Topography of efferent vagal innervation of the rat gastrointestinal tract. Am J Physiol 260(1 Pt 2): R200-207.

Brierley, S. M., R. C. Jones, G. F. Gebhart and L. A. Blackshaw (2004). Splanchnic and pelvic mechanosensory afferents signal different qualities of colonic stimuli in mice. Gastroenterology 127: 166–178.

Browne, T. J., K. M. Smith, M. A. Gradwell, J. A. Iredale, C. V. Dayas, R. J. Callister, . . . B. A. Graham (2021). Spinoparabrachial projection neurons form distinct classes in the mouse dorsal horn. Pain 162(7): 1977–1994.

Cameron, D., E. Polgar, M. Gutierrez-Mecinas, M. Gomez-Lima, M. Watanabe and A. J. Todd (2015). The organisation of spinoparabrachial neurons in the mouse. Pain 156(10): 2061-2071.

Cedarbaum, J. M. and G. K. Aghajanian (1978). Afferent projections to the rat locus coeruleus as determined by a retrograde tracing technique. J Comp Neurol 178(1): 1–16.

Chiang, M. C., E. K. Nguyen, M. Canto-Bustos, A. E. Papale, A. M. Oswald and S. E. Ross (2020). Divergent Neural Pathways Emanating from the Lateral Parabrachial Nucleus Mediate Distinct Components of the Pain Response. Neuron 106(6): 927–939 e925.

Choi, S., J. Hachisuka, M. A. Brett, A. R. Magee, Y. Omori, N. U. Iqbal, . . . D. D. Ginty (2020). Parallel ascending spinal pathways for affective touch and pain. Nature 587(7833): 258–263.

Christianson, J. A. and G. F. Gebhart (2007). Assessment of colon sensitivity by luminal distension in mice. Nat Protoc 2(10): 2624–2631.

Christianson, J. A., R. J. Traub and B. M. Davis (2006). Differences in spinal distribution and neurochemical phenotype of colonic afferents in mouse and rat. J Comp Neurol 494(2): 246–259.

Churchill, M. J., M. A. Cantu, E. A. Kasanga, C. Moore, M. F. Salvatore and C. K. Meshul (2019). Glatiramer Acetate Reverses Motor Dysfunction and the Decrease in Tyrosine Hydroxylase Levels in a Mouse Model of Parkinson’s Disease. Neuroscience 414: 8–27.

Ding, Y. Q., H. X. Zheng, L. W. Gong, Y. Lu, H. Zhao and B. Z. Qin (1997). Direct projections from the lumbosacral spinal cord to Barrington’s nucleus in the rat: a special reference to micturition reflex. J Comp Neurol 389(1): 149–160.

Garcia-Luna, C., G. Sanchez-Watts, M. Arnold, G. de Lartigue, N. DeWalt, W. Langhans and A. G. Watts (2021). The Medullary Targets of Neurally Conveyed Sensory Information from the Rat Hepatic Portal and Superior Mesenteric Veins. eNeuro 8(1).

Gu, X., Y. Z. Zhang, J. J. O’Malley, C. C. De Preter, M. Penzo and M. A. Hoon (2023). Neurons in the caudal ventrolateral medulla mediate descending pain control. Nat Neurosci 26(4): 594–605.

Harrington, A. M. (2023). Translating Colonic Sensory Afferent Peripheral Mechanosensitivity into the Spinal Cord Dorsal Horn. Visceral Pain. S. M. Brierley and N. J. Spencer, Springer, Cham: 161–181.

Harrington, A. M., S. G. Caraballo, J. E. Maddern, L. Grundy, J. Castro and S. M. Brierley (2019). Colonic afferent input and dorsal horn neuron activation differs between the thoracolumbar and lumbosacral spinal cord. Am J Physiol Gastrointest Liver Physiol 317(3): G285–G303.

He, Z. G., Q. Wang, R. S. Xie, Y. S. Li, Q. X. Hong and H. B. Xiang (2018). Neuroanatomical autonomic substrates of brainstem-gut circuitry identified using transsynaptic tract-tracing with pseudorabies virus recombinants. Am J Clin Exp Immunol 7(2): 16–24.

Hingorani, M., A. M. L. Viviani, J. E. Sanfilippo and S. Janusonis (2022). High-resolution spatiotemporal analysis of single serotonergic axons in an in vitro system. Front Neurosci 16: 994735.

Iwasaki, K., H. Komiya, M. Kakizaki, C. Miyoshi, M. Abe, K. Sakimura, . . . M. Yanagisawa (2018). Ablation of Central Serotonergic Neurons Decreased REM Sleep and Attenuated Arousal Response. Front Neurosci 12: 535.

Kawatani, M., W. C. de Groat, K. Itoi, K. Uchida, K. Sakimura, A. Yamanaka, . . . M. Kawatani (2021). Downstream projection of Barrington’s nucleus to the spinal cord in mice. J Neurophysiol 126(6): 1959–1977.

Keay, K. A., K. Feil, B. D. Gordon, H. Herbert and R. Bandler (1997). Spinal afferents to functionally distinct periaqueductal gray columns in the rat: an anterograde and retrograde tracing study. J Comp Neurol 385(2): 207–229.

Kyloh, M. A., T. J. Hibberd, J. Castro, A. M. Harrington, L. Travis, K. N. Dodds, . . . N. J. Spencer (2022). Disengaging spinal afferent nerve communication with the brain in live mice. Commun Biol 5(1): 915.

Lein, E. S., M. J. Hawrylycz, N. Ao, M. Ayres, A. Bensinger, A. Bernard, . . . A. R. Jones (2007). Genome-wide atlas of gene expression in the adult mouse brain. Nature 445(7124): 168–176.

Lima, D., J. A. Mendes-Ribeiro and A. Coimbra (1991). The spino-latero-reticular system of the rat: projections from the superficial dorsal horn and structural characterization of marginal neurons involved. Neuroscience 45(1): 137–152.

Liu, L., M. Tsuruoka, M. Maeda, B. Hayashi and T. Inoue (2007). Coeruleospinal inhibition of visceral nociceptive processing in the rat spinal cord. Neurosci Lett 426(3): 139–144.

Liu, L., M. Tsuruoka, M. Maeda, B. Hayashi, X. Wang and T. Inoue (2008). Descending modulation of visceral nociceptive transmission from the locus coeruleus/subcoeruleus in the rat. Brain Res Bull 76(6): 616–625.

Lubejko, S. T., G. Livrizzi, S. A. Buczynski, J. Patel, J. C. Yung, T. L. Yaksh and M. R. Banghart (2024). Inputs to the locus coeruleus from the periaqueductal gray and rostroventral medulla shape opioid-mediated descending pain modulation. Sci Adv 10(17): eadj9581.

Lyubashina, O. A., I. B. Sivachenko and A. A. Mikhalkin (2022). Impaired visceral pain-related functions of the midbrain periaqueductal gray in rats with colitis. Brain Res Bull 182: 12–25.

Lyubashina, O. A., I. B. Sivachenko and A. Y. Sokolov (2019). Differential responses of neurons in the rat caudal ventrolateral medulla to visceral and somatic noxious stimuli and their alterations in colitis. Brain Res Bull 152: 299–310.

Martins, I. and I. Tavares (2017). Reticular Formation and Pain: The Past and the Future. Front Neuroanat 11: 51.

McGovern, A. E., N. Davis-Poynter, M. J. Farrell and S. B. Mazzone (2012). Transneuronal tracing of airways-related sensory circuitry using herpes simplex virus 1, strain H129. Neuroscience 207: 148–166.

McGovern, A. E., N. Davis-Poynter, J. Rakoczy, S. Phipps, D. G. Simmons and S. B. Mazzone (2012). Anterograde neuronal circuit tracing using a genetically modified herpes simplex virus expressing EGFP. J Neurosci Methods 209(1): 158–167.

McGovern, A. E., N. Davis-Poynter, S. K. Yang, D. G. Simmons, M. J. Farrell and S. B. Mazzone (2015). Evidence for multiple sensory circuits in the brain arising from the respiratory system: an anterograde viral tract tracing study in rodents. Brain Struct Funct 220(6): 3683–3699.

Miyaji, M., R. L. Kortum, R. Surana, W. Li, K. D. Woolard, R. M. Simpson, . . . C. L. Sommers (2009). Genetic evidence for the role of Erk activation in a lymphoproliferative disease of mice. Proc Natl Acad Sci U S A 106(34): 14502–14507.

Monnikes, H., J. Ruter, M. Konig, C. Grote, P. Kobelt, B. F. Klapp, . . . J. J. Tebbe (2003). Differential induction of c-fos expression in brain nuclei by noxious and non-noxious colonic distension: role of afferent C-fibers and 5-HT3 receptors. Brain Res 966(2): 253–264.

Monnikes, H., B. G. Schmidt, J. Tebbe, C. Bauer and Y. Tache (1994). Microinfusion of corticotropin releasing factor into the locus coeruleus/subcoeruleus nuclei stimulates colonic motor function in rats. Brain Res 644(1): 101–108.

Murphy, A. Z., S. K. Suckow, M. Johns and R. J. Traub (2009). Sex differences in the activation of the spinoparabrachial circuit by visceral pain. Physiol Behav 97(2): 205–212.

Naitou, K., H. Nakamori, K. Horii, K. Kato, Y. Horii, H. Shimaoka, . . . Y. Shimizu (2018). Descending monoaminergic pathways projecting to the spinal defecation center enhance colorectal motility in rats. Am J Physiol Gastrointest Liver Physiol 315(4): G631–G637.

Nakamori, H., K. Naitou, Y. Horii, H. Shimaoka, K. Horii, H. Sakai, . . . Y. Shimizu (2018). Medullary raphe nuclei activate the lumbosacral defecation center through the descending serotonergic pathway to regulate colorectal motility in rats. Am J Physiol Gastrointest Liver Physiol 314(3): G341–G348.

Nakamori, H., K. Naitou, Y. Horii, H. Shimaoka, K. Horii, H. Sakai, . . . Y. Shimizu (2019). Roles of the noradrenergic nucleus locus coeruleus and dopaminergic nucleus A11 region as supraspinal defecation centers in rats. Am J Physiol Gastrointest Liver Physiol 317(4): G545–G555.

Ness, T. J., K. A. Follett, J. Piper and B. A. Dirks (1998). Characterization of neurons in the area of the medullary lateral reticular nucleus responsive to noxious visceral and cutaneous stimuli. Brain Res 802(1-2): 163–174.

Ness, T. J. and G. F. Gebhart (1988). Colorectal distension as a noxious visceral stimulus: physiologic and pharmacologic characterization of pseudaffective reflexes in the rat. Brain Res 450(1-2): 153–169.

Nishida, K., S. Matsumura and T. Kobayashi (2022). Involvement of Brn3a-positive spinal dorsal horn neurons in the transmission of visceral pain in inflammatory bowel disease model mice. Front Pain Res (Lausanne) 3: 979038.

Okada, K., K. Murase and K. Kawakita (1999). Effects of electrical stimulation of thalamic nucleus submedius and periaqueductal gray on the visceral nociceptive responses of spinal dorsal horn neurons in the rat. Brain Res 834(1-2): 112–121.

Osman, S., A. Tashtush, D. E. Reed and A. E. Lomax (2023). Analysis of the spinal and vagal afferent innervation of the mouse colon using neuronal retrograde tracers. Cell Tissue Res 392(3): 659–670.

Parker, C. G., M. J. Dailey, H. Phillips and E. A. Davis (2020). Central sensory-motor crosstalk in the neural gut-brain axis. Auton Neurosci 225: 102656.

Pavcovich, L. A., M. Yang, R. R. Miselis and R. J. Valentino (1998). Novel role for the pontine micturition center, Barrington’s nucleus: evidence for coordination of colonic and forebrain activity. Brain Res 784(1-2): 355–361.

Paxinos, G. and K. Franklin (2001). The Mouse Brain in Stereotaxic Coordinates, Second Edition. California, Academic Press.

Peirs, C., R. Dallel and A. J. Todd (2020). Recent advances in our understanding of the organization of dorsal horn neuron populations and their contribution to cutaneous mechanical allodynia. J Neural Transm (Vienna) 127(4): 505–525.

Peng, B., Y. Jiao, Y. Zhang, S. Li, S. Chen, S. Xu, . . . W. Yu (2023). Bulbospinal nociceptive ON and OFF cells related neural circuits and transmitters. Front Pharmacol 14: 1159753.

Polgar, E., L. L. Wright and A. J. Todd (2010). A quantitative study of brainstem projections from lamina I neurons in the cervical and lumbar enlargement of the rat. Brain Res 1308(5): 58–67.

Qi, L., S. H. Lin and Q. Ma (2022). Spinal VGLUT3 lineage neurons drive visceral mechanical allodynia but not sensitized visceromotor reflexes. Neuron.

Rinaman, L. and G. Schwartz (2004). Anterograde transneuronal viral tracing of central viscerosensory pathways in rats. J Neurosci 24(11): 2782–2786.

Robinson, D. R., P. A. McNaughton, M. L. Evans and G. A. Hicks (2004). Characterization of the primary spinal afferent innervation of the mouse colon using retrograde labelling. Neurogastroenterol Motil 16(1): 113–124.

Roeder, Z., Q. Chen, S. Davis, J. D. Carlson, D. Tupone and M. M. Heinricher (2016). Parabrachial complex links pain transmission to descending pain modulation. Pain 157(12): 2697–2708.

Rouzade-Dominguez, M. L., R. Miselis and R. J. Valentino (2003). Central representation of bladder and colon revealed by dual transsynaptic tracing in the rat: substrates for pelvic visceral coordination. Eur J Neurosci 18(12): 3311–3324.

Rouzade-Dominguez, M. L., L. Pernar, S. Beck and R. J. Valentino (2003). Convergent responses of Barrington’s nucleus neurons to pelvic visceral stimuli in the rat: a juxtacellular labelling study. Eur J Neurosci 18(12): 3325–3334.

Samuels, E. R. and E. Szabadi (2008). Functional neuroanatomy of the noradrenergic locus coeruleus: its roles in the regulation of arousal and autonomic function part II: physiological and pharmacological manipulations and pathological alterations of locus coeruleus activity in humans. Curr Neuropharmacol 6(3): 254–285.

Sikandar, S. and A. H. Dickenson (2012). Visceral pain: the ins and outs, the ups and downs. Curr Opin Support Palliat Care 6(1): 17–26.

Spencer, N. J., M. A. Kyloh, L. Travis and T. J. Hibberd (2024). Identification of vagal afferent nerve endings in the mouse colon and their spatial relationship with enterochromaffin cells. Cell Tissue Res 396(3): 313–327.

Tao, J., J. N. Campbell, L. T. Tsai, C. Wu, S. D. Liberles and B. B. Lowell (2021). Highly selective brain-to-gut communication via genetically defined vagus neurons. Neuron 109(13): 2106–2115 e2104.

Tavares, I. and D. Lima (2002). The caudal ventrolateral medulla as an important inhibitory modulator of pain transmission in the spinal cord. J Pain 3(5): 337–346.

Todd, A. J. (2010). Neuronal circuitry for pain processing in the dorsal horn. Nat Rev Neurosci 11(12): 823–836.

Valentino, R. J., M. Kosboth, M. Colflesh and R. R. Miselis (2000). Transneuronal labeling from the rat distal colon: anatomic evidence for regulation of distal colon function by a pontine corticotropin-releasing factor system. J Comp Neurol 417(4): 399–414.

Vizzard, M. A., M. Brisson and W. C. de Groat (2000). Transneuronal labeling of neurons in the adult rat central nervous system following inoculation of pseudorabies virus into the colon. Cell Tissue Res 299(1): 9–26.

Wang, C. C., W. D. Willis and K. N. Westlund (1999). Ascending projections from the area around the spinal cord central canal: A Phaseolus vulgaris leucoagglutinin study in rats. J Comp Neurol 415(3): 341–367.

Wang, F. B. and T. L. Powley (2000). Topographic inventories of vagal afferents in gastrointestinal muscle. J Comp Neurol 421(3): 302–324.

Wang, Q., S. G. Caraballo, G. Rychkov, A. E. McGovern, S. B. Mazzone, S. M. Brierley and A. M. Harrington (2024). Comparative localization of colorectal sensory afferent central projections in the mouse spinal cord dorsal horn and caudal medulla dorsal vagal complex. J Comp Neurol 532(2): e25546.

Wercberger, R. and A. I. Basbaum (2019). Spinal cord projection neurons: a superficial, and also deep, analysis. Curr Opin Physiol 11: 109–115.

Westlund, K. N. (2000). Visceral nociception. Curr Rev Pain 4(6): 478–487.

Wojaczynski, G. J., E. A. Engel, K. E. Steren, L. W. Enquist and J. Patrick Card (2015). The neuroinvasive profiles of H129 (herpes simplex virus type 1) recombinants with putative anterograde-only transneuronal spread properties. Brain Struct Funct 220(3): 1395–1420.

